# Nearest Neighbor Parameters for Estimating RNA Folding Stability with *In Vivo*-like Conditions

**DOI:** 10.64898/2026.09.04.749202

**Authors:** Olivia M. Hiltke, Elzbieta Kierzek, Martina Prochota, Marta Rachwalak, Megan Miaro, Thandolwethu Shabangu, Sylwia Gorczynska, Ryszard Kierzek, David H. Mathews

## Abstract

RNAs regulate gene expression and cellular processes, often relying on specific conformations for function. RNA folding is hierarchical and sequence-dependent, with nearest-neighbor thermodynamic models commonly used to predict secondary structure. Current models were developed using optical melting experiments in 1 M NaCl, which does not represent the cellular environment. To address this, we developed a new model in Advanced Dulbecco’s Modified Eagle Medium (Adv. DMEM), which mimics mammalian extracellular ionic composition. This *in vivo*-like model provides RNA folding parameters for helical base stacks and loop motifs. Optical melting experiments revealed helical stacks, particularly tandem G-U pairs, are less stabilizing in Adv. DMEM. Loop parameters were generally destabilizing but highly dependent on both sequence and loop type, with internal loops displaying idiosyncratic behavior. Structure prediction benchmarking revealed minimal differences overall, except for tRNAs, which showed improved prediction reliability and enhanced cloverleaf stability. Notably, tRNAs lack internal loops, suggesting further studies in Adv. DMEM could refine secondary structure predictions. This *in vivo*-like parameter set is included in the RNAstructure software package. Grounding these parameters in a physiologically relevant environment, we improve the biological relevance of RNA secondary structure predictions and establish a foundation for studying RNA folding under *in vivo* conditions.

**Graphical Abstract:** 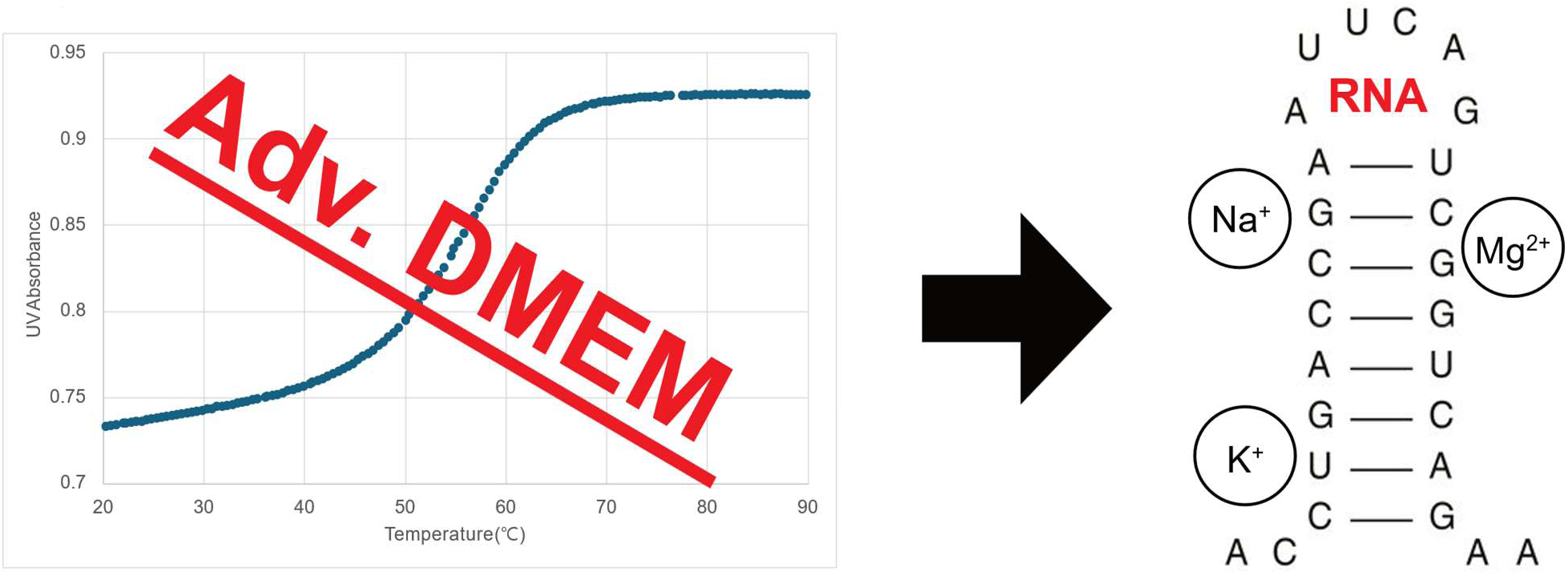

## Introduction

In the past, RNA was mainly recognized for its role as mRNA in the translation of DNA into proteins. Today, RNA is recognized as a central regulator of cellular function. Although over 80% of the human genome is transcribed into RNA, less than 3% encodes proteins [1,2]; this pervasive transcription suggests the vast majority of RNA transcripts are non-coding RNAs (ncRNAs) that function without being translated into proteins. These ncRNAs are essential in many cellular processes including post-transcriptional regulation [3–5], catalysis [3,4,6], protein trafficking [7], and guiding site-specific RNA modifications [8]. Despite the recent rapid pace of discovery, genome-wide transcription suggests that many ncRNA species and mechanisms remain unidentified [9]. Discovering and characterizing these ncRNAs is essential not only for advancing basic understanding of RNA biology, but also for identifying novel targets for therapeutic intervention and biotechnology applications.

The functions of ncRNAs depend on folding into specific structures [10,11]. Accurate structure prediction is therefore critical for understanding the roles of RNA molecules. RNA structure is hierarchical, with canonical base pairs forming faster and being more thermostable than tertiary structure contacts [12–15]. RNA secondary structure is defined by the canonical base pairings that arrange the strands into A-form helices, which are connected by characteristic loop motifs including hairpin loops, internal loops, bulge loops, multibranch loops, and exterior loops [16,17]. Modeling secondary structure provides a characterization of the RNA family and provides a framework for modeling three-dimensional structure [18,19], which in turn offers insight into the RNA’s function.

The conformational folding stability of an RNA secondary structure is quantified as a standard state Gibbs free energy change at 37 °C (ΔG°_37)_, where the conformation with the lowest folding free energy change is the most thermodynamically stable and, therefore, the most populated structure at equilibrium [20–22]. Thermodynamic models are used to estimate the ΔG°_37_ associated with base pairing and loop motifs. The current approach is the nearest-neighbor model, which assumes that the stability of each motif depends on only the sequence of that motif and the most adjacent base pairs [23]. The total ΔG°_37_ of a structure is the sum of these parameters (Figure 1).

**Figure 1.**
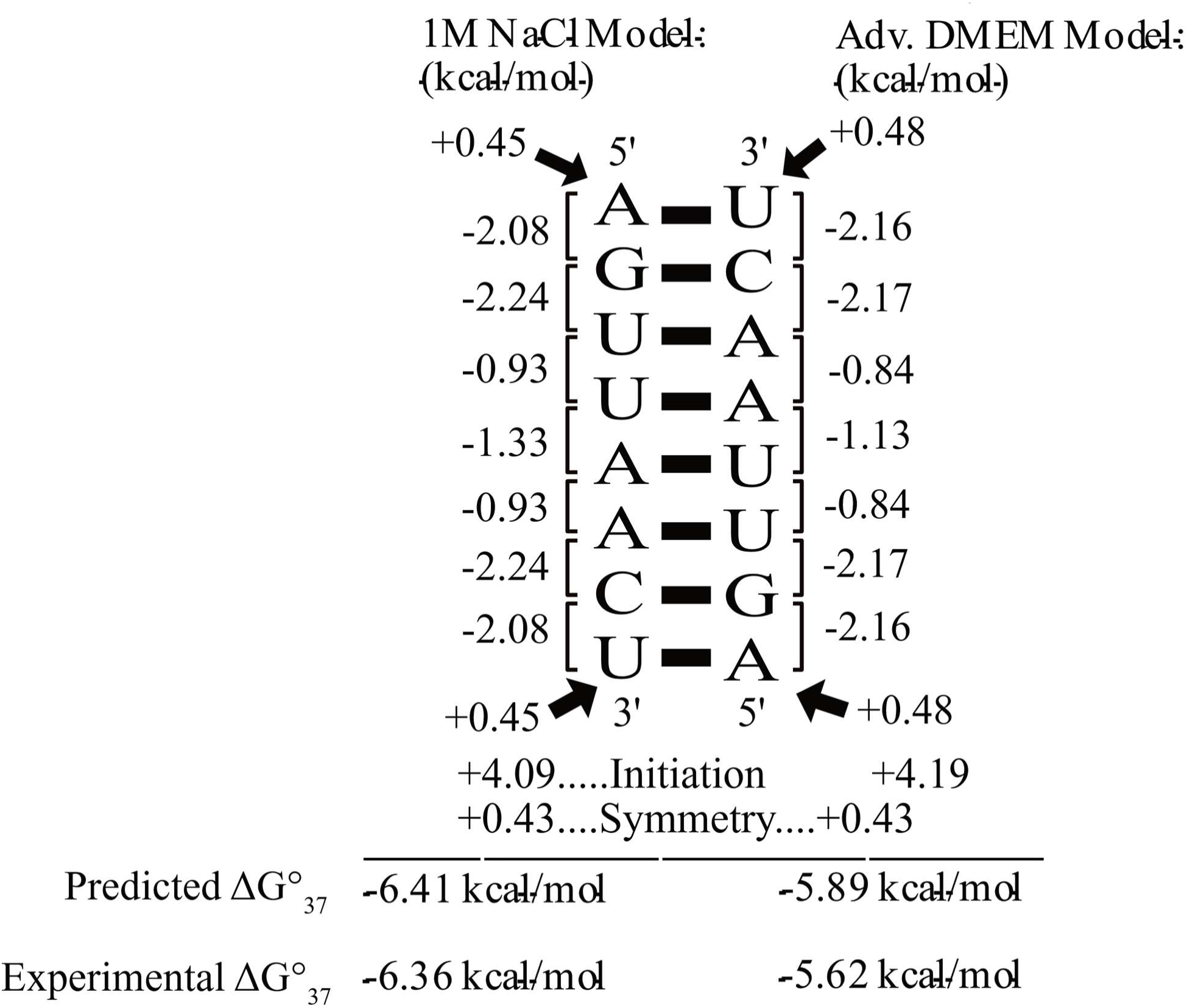
Comparison of traditional 1 M Na^+^ and *in vivo-like* Nearest Neighbor Calculations for the Folding Free Energy Change of an RNA Helix. The left column shows the calculation and experiment in 1 M Na^+^ and the right column shows the calculation and experiment in Adv. DMEM. The helical stack parameters are dependent on the sequence of the base pair and its neighboring pair. The arrows indicate A-U pairs at the helix termini that receive a penalty term. The helix initiation penalty compensates for the entropy loss when two individual RNA strands form a duplex. The symmetry correction penalty applies when a duplex is formed between identical (self-complementary) strands. The total estimated free energy change of an RNA sequence is the sum of all the applicable nearest-neighbor parameter terms.

The nearest-neighbor parameters were developed to reproduce experimentally measured stabilities from RNA sequences using UV optical melting experiments [21,24,25]. The current set of parameters, called Turner 2004, has terms for canonical helical pairs and for loops, which creates the foundation for tools such as NUPACK, RNAstructure, UNAFold, and the Vienna Package, which use dynamic programming algorithms to predict the lowest free energy structure of an RNA sequence [23,26–29]. These tools are widely used in the field to investigate RNA function and design RNA-based therapeutics [22]. Therefore, improving the accuracy of RNA structure prediction not only deepens the understanding of RNA biology but also advances applications in medicine, biotechnology, and evolutionary biology.

The Turner 2004 model was developed from optical melting experiments performed in 1 M NaCl [24,25,30]. This media composition has been used for optical melting measurements since the early 1970s and was chosen to provide strong electrostatic screening for the negatively charged RNA backbone while avoiding the inclusion of Mg²⁺, which can catalyze RNA cleavage at high temperatures [23–25,27,31]. At the time, the synthesis and purification of RNA were labor-intensive and costly. Therefore, preserving samples during melting experiments was a priority. Over time, the use of 1 M NaCl became the standardized framework for nucleic acid thermodynamics, enabling direct comparison across datasets. However, this high sodium, Mg²⁺-free environment does not reflect the complex ionic conditions found in cells, where physiological concentrations of monovalent and divalent cations can impose substantial influence on RNA stability and conformation. Consequently, parameters derived under 1 M NaCl may not fully capture how RNA structures form and function *in vivo* [32,33].

The *in vitro* environment used for RNA optical melting studies only partially relates to the conditions present in the cells. Three prior publications focused on the stabilities of RNA under *in vivo*-like conditions to get the thermodynamic parameters necessary to predict folding of RNA in the cells [33–35]. Adams and Znosko and Ghosh et al. focused on the influence of crowding conditions on folding RNA and used 20% and 40%, respectively, polyethylene glycol-200 (PEG-200) to mimic a cellular environment [34,35]. Sieg et al. used an Eco80 artificial cytoplasm as the medium to measure the thermodynamic stability of RNA [33]. In these studies, the thermodynamic measurements were limited to RNA helices only and therefore did not provide information for loops. The loop terms in the Turner 2004 rules are also important for precise secondary structure prediction [36].

The problem of determining the folding stability of RNA in cellular conditions is challenging because there is no universal cellular condition. Dependent on the type of cells and organelles within the cell, the composition and concentration of ions, carbohydrates, proteins, metabolites, nucleic acids, amino acids, and other components vary [37]. For example, in *Escherichia coli, Saccharomyces cerevisiae*, and mammalian cells, the concentrations of potassium cations are 30-300, 300, and 100 mM, whereas the concentrations of sodium cations are 10, 30, and 10 mM, respectively. In mammalian cells, intracellular fluid contains mostly potassium cations, whereas intercellular fluid contains mostly sodium cations. Also, intracellular pH values vary depending on cellular organelles in the range between 4.5 and 8.0 [37,38].

Here, we report nearest-neighbor parameters for canonical helical stacks, terminal mismatches, dangling ends, bulge loops, and hairpin loops under *in vivo*-like conditions. A set of 193 optical melting experiments were performed in Advanced Dulbecco’s Modified Eagle Medium (Adv. DMEM), a mammalian cell culture medium supplemented a mammalian cell culture medium with nutrients and physiologically relevant ion concentrations (Table 1). By parameterizing to these conditions, this new model addresses a key limitation of the current Turner 2004 rules and has the potential to improve the biological relevance of RNA secondary structure predictions. We found, however, that we could not parameterize internal loops using a limited set of 16 experiments because their stabilities in Adv. DMEM do not correlate to those in 1 M Na^+^. The new *in vivo-*like parameters were incorporated into the RNAstructure software package and are freely available. We found that they significantly improve the accuracy of structure prediction for tRNAs, an RNA family that almost completely lacks internal loops. The structure prediction accuracies for the other RNA families were generally not improved.

**Table 1.** Composition of Advanced Dulbecco’s Modified Eagle Medium (Adv. DMEM). Solution was prepared using the D2902 formulation (Sigma-Aldrich), a powdered low-glucose DMEM containing inorganic salts, amino acids, vitamins, and other essential nutrients for mammalian cell culture [71] (https://www.sigmaaldrich.com/deepweb/assets/sigmaaldrich/product/documents/418/100/d2902for.pdf). Concentrations are reported as grams per liter (g/L) of final 1× medium. Sodium bicarbonate (3.7 g/L) was added separately during medium preparation to provide buffering capacity and maintain physiological pH

| Component | Concentration (g/L) |
| --- | --- |
| Calcium chloride ($\text{CaCl}_2$) | 0.2000 |
| Ferric nitrate nonahydrate [$\text{Fe}(\text{NO}_3)_3 \cdot 9\text{H}_2\text{O}$] | 0.0001 |
| Magnesium sulfate ($\text{MgSO}_4$) | 0.0977 |
| Potassium chloride (KCl) | 0.4000 |
| Sodium chloride (NaCl) | 6.4000 |
| Sodium phosphate monobasic ($\text{NaH}_2\text{PO}_4$) | 0.1090 |
| L – Arginine · HCl | 0.0840 |
| L – Cystine · 2HCl | 0.0626 |
| L-Glutamine | 0.5840 |
| Glycine | 0.0300 |
| L-Histidine · HCl · $\text{H}_2\text{O}$ | 0.0420 |
| L-Isoleucine | 0.1050 |
| L-Leucine | 0.1050 |
| L-Lysine · HCl | 0.1460 |
| L-Methionine | 0.0300 |
| L-Phenylalanine | 0.0660 |
| L-Serine | 0.0420 |
| L-Threonine | 0.0950 |
| L-Tryptophan | 0.0160 |
| L-Tyrosine disodium salt dihydrate | 0.1038 |
| L-Valine | 0.0940 |
| Choline chloride | 0.0040 |
| Folic acid | 0.0040 |
| myo-Inositol | 0.0072 |
| Niacinamide | 0.0040 |
| D-Pantothenic acid · $\frac{1}{2}\text{Ca}$ | 0.0040 |
| Pyridoxal · HCl | 0.0040 |
| Riboflavin | 0.0004 |
| Thiamine · HCl | 0.0040 |
| D-Glucose | 1.0000 |
| Sodium pyruvate | 0.1100 |
| Sodium bicarbonate ($\text{NaHCO}_3$) | 3.7000 |

## Material and Methods

### Oligonucleotides Synthesis

Oligonucleotides were synthesized on a BioAutomation MerMade12 DNA/RNA synthesizer using β-cyanoethyl phosphoramidite chemistry [39] and commercially available phosphoramidites (ChemGenes, GenePharma). The synthesis was performed using the standard RNA protocol. For deprotection, oligoribonucleotides were treated with a mixture of 30% aqueous ammonia/ethanol (3/1 v/v) for 16 h at 55 ^°^C. Silyl protecting groups were cleaved by treatment with trimethylamine trihydrofluoride. Deprotected oligonucleotides were purified by silica gel thin layer chromatography (TLC) in 1-propanol/aqueous ammonia/water (55/35/10 v/v/v) as described in detail previously [24,40].

### UV Melting Experiments

The thermodynamic measurements were performed for nine various concentrations of RNA duplex in the range 0.1 mM to 1 µM on a JASCO V-650 UV/Vis spectrophotometer in Adv. DMEM solution (Dulbecco’s Modified Eagle’s Medium -low glucose with 1000 mg/L glucose and L-glutamine, without sodium bicarbonate and phenol red, Sigma catalog number D2902) in quartz cells with pathlengths 0.1, 0.5, and 1.0 cm. To make 10 ml of Adv. DMEM solution, 100 mg of powder was dissolved in 10 ml of ultrapure (MQ) water. Oligonucleotide single-strand concentrations were calculated from the absorbance above 80 °C, and single-strand extinction coefficients were approximated by a nearest-neighbor model [41,42].

Absorbance vs. temperature melting curves were measured at 260 nm with a heating rate of 1 °C/min from 0 to about 90 °C on a JASCO V-650 spectrophotometer with a thermoprogrammer, where a reference was Adv. DMEM solution in 1.0 cm quartz cell (room temperature). Before performing UV-melting of RNA duplexes, the solution of Adv. DMEM alone was melted in 0.1, 0.5, and 1.0 cm cells. In that case, as a reference, a 1.0 cm quartz cell containing Adv. DMEM solution was used (room temperature). The presence of some proteins in Adv. DMEM results in absorbance at 260 nm, specific to a particular cell pathlength and temperature. This UV-melting of Adv. DMEM solution was performed 5 times and averaged to obtain accurate results. The thermodynamic measurements in Adv. DMEM were performed for nine various concentrations of RNA as described above, and each of the nine samples was prepared from a new RNA and Adv. DMEM solution to avoid the presence of denatured proteins present in Adv. DMEM solution. To correct the effect of proteins, the “reference” Adv. DMEM absorbance values specific to the quartz cuvette pathlength were subtracted from the “cell pathlength”-specific RNA UV absorbance at all temperatures. The melting curves were then analyzed and the thermodynamic parameters were calculated from a two-state model with the program MeltWin 3.5 [43]. For most sequences, the ΔH° derived from TM^-1^ vs. ln(C_T_/4) plots is within 15% of that derived from averaging the fits to individual melting curves, as expected if the two-state model is reasonable.

### Helical Stack Parameters

Helical stack parameters, including both Watson-Crick-Franklin (WCF) base pairs and G-U base pairs were fit by non-error-weighted linear regression. The folding stability measurements (ΔG°₃₇ measurements from TM^-1^ vs. ln(C_T_/4) plots) of 42 RNA helices containing WCF base pairs in Adv. DMEM were used to fit the 10 stack parameters for WCF base stacking combinations, as well as additional parameters for helix initiation and A-U terminal end (Figure 2A, Supplementary Tables 1 and 2). A custom Python program was written to perform the regression. It used the *statsmodel* and *Numpy* libraries [44,45].

**Figure 2.**
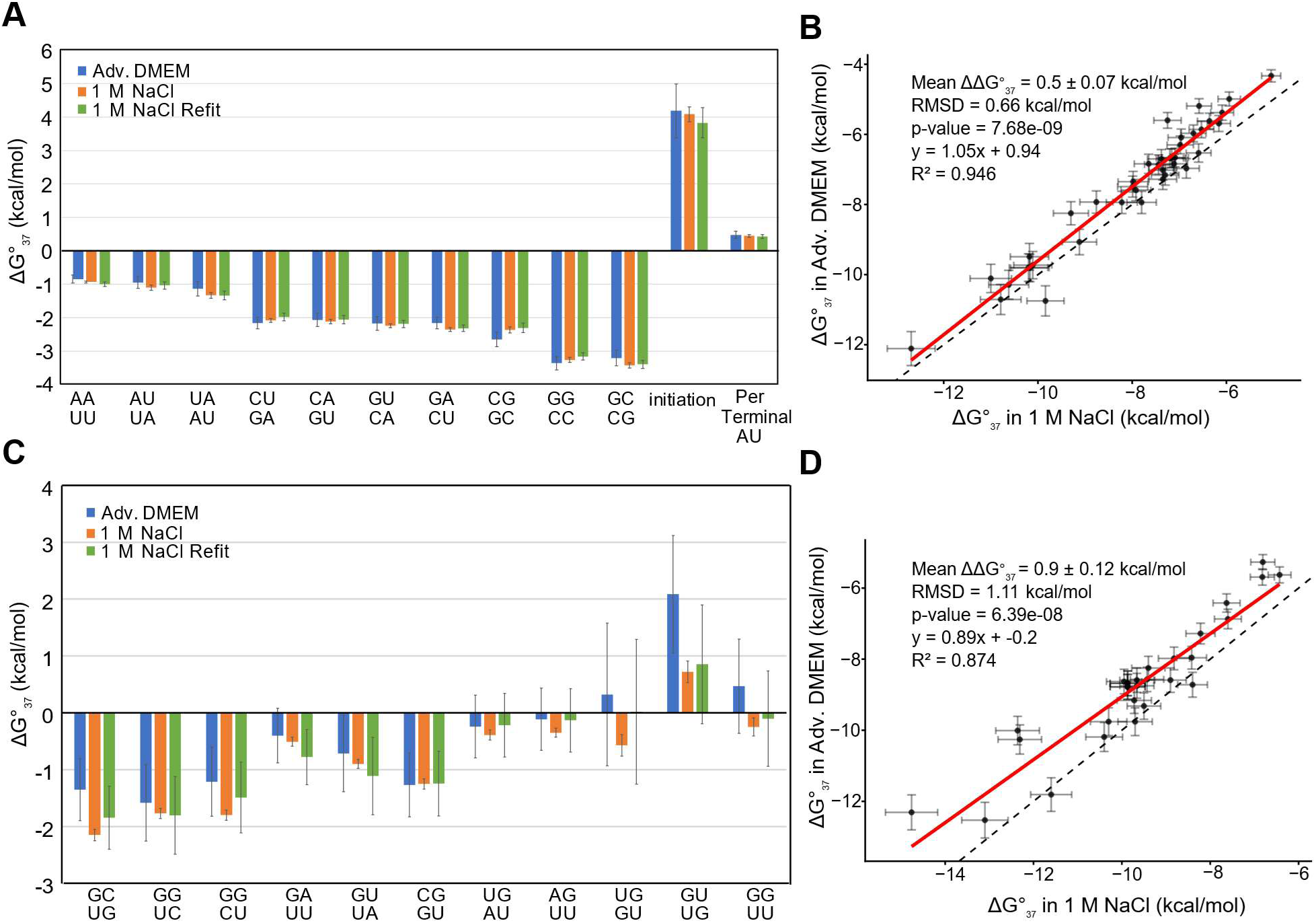
Helical Stacks Thermodynamic Behavior in Adv. DMEM Versus 1 M NaCl. **(A)** Nearest neighbor parameters for Watson-Crick-Franklin (WCF) base pair stacks, helix initiation, and A-U terminal ends were determined from folding stability of WCF helices measured via UV optical melting in Adv. DMEM of 42 helices (blue) and standard Turner 2004 parameters derived in 1 M NaCl with 90 helices (orange) [24]. WCF nearest-neighbor parameters were recalculated for 40 helices measured in the Adv. DMEM dataset (Supplementary Table 1) using the folding free energy measured in 1 M NaCl (green) to facilitate a direct comparison to the Adv. DMEM parameters [24]. The error bars represent the uncertainty estimate using the standard error of the regression. **(B)** ΔG°_37_ of WCF-containing helices measured via UV optical melting in Adv. DMEM versus 1 M NaCl. Points and error bars represent the ΔG°_37_ determined using Tm^-1^ vs. ln(C_T_) fits and experimental error [24]. The R^2^ represents the coefficient of determination of the linear model represented graphically with the red line. The dashed line represents the diagonal. The two-tailed paired t-test between the folding stabilities of the helices measured in the two conditions was performed using the t.test function in R. **(C)** Nearest neighbor parameters for G-U base pair stacks were determined from folding stability of G-U-containing helices measured via UV optical melting in Advanced DMEM of 23 helices (blue) and revised G-U parameters derived in 1 M NaCl of 39 helices (orange) [30]. G-U nearest-neighbor parameters were recalculated for the 23 helices measured in the Adv. DMEM dataset (Supplementary Table 3) using the folding free energy measured in 1 M NaCl (green). The error bars represent the uncertainty estimate using the standard error of the regression. **(D)** ΔG°_37_ of G-U-containing helices measured via UV optical melting in Adv. DMEM versus 1 M NaCl. Points and error bars represent the ΔG°_37_ and experimental error, respectively, determined Tm^-1^ vs. ln(C_T_) fits [30]. The R^2^ represents the coefficient of determination of the linear model presented by the figure and represented graphically with the red line. The dashed line represents the diagonal. The two-tailed paired t-test between the folding stabilities of the helices measured in the two conditions was performed using the t.test function in R.

The 11 G-U stack parameters were subsequently fit in a second linear regression using a dataset of 23 G-U-containing duplexes (Figure 2C, Supplementary Tables 3 and 4), following the practice used for 1 M Na^+^ parameter derivation [25,30]. WCF stacks, helix initiation and A-U terminal terms were subtracted from the duplex ΔG°₃₇ to isolate G-U contributions for the linear regression fit and uncertainty estimates. We determined that no terminal stability penalties were needed for terminal G-U pairs, similar to a revision of G-U parameters in 1 M Na^+^ that found the penalty for terminal G-U pairs to be unnecessary [30].

Self-complementary duplexes were corrected for symmetry prior to linear regression by subtracting 0.43 kcal/mol from the measured free energy change of the RNA sequence. This penalty accounts for reduced entropy, as the two strands are indistinguishable [24].

The standard error of the linear regression was used to estimate helical parameter uncertainty [24]. The R^2^ for the helical parameter fit was 0.856 (Figure 3).

**Figure 3.**
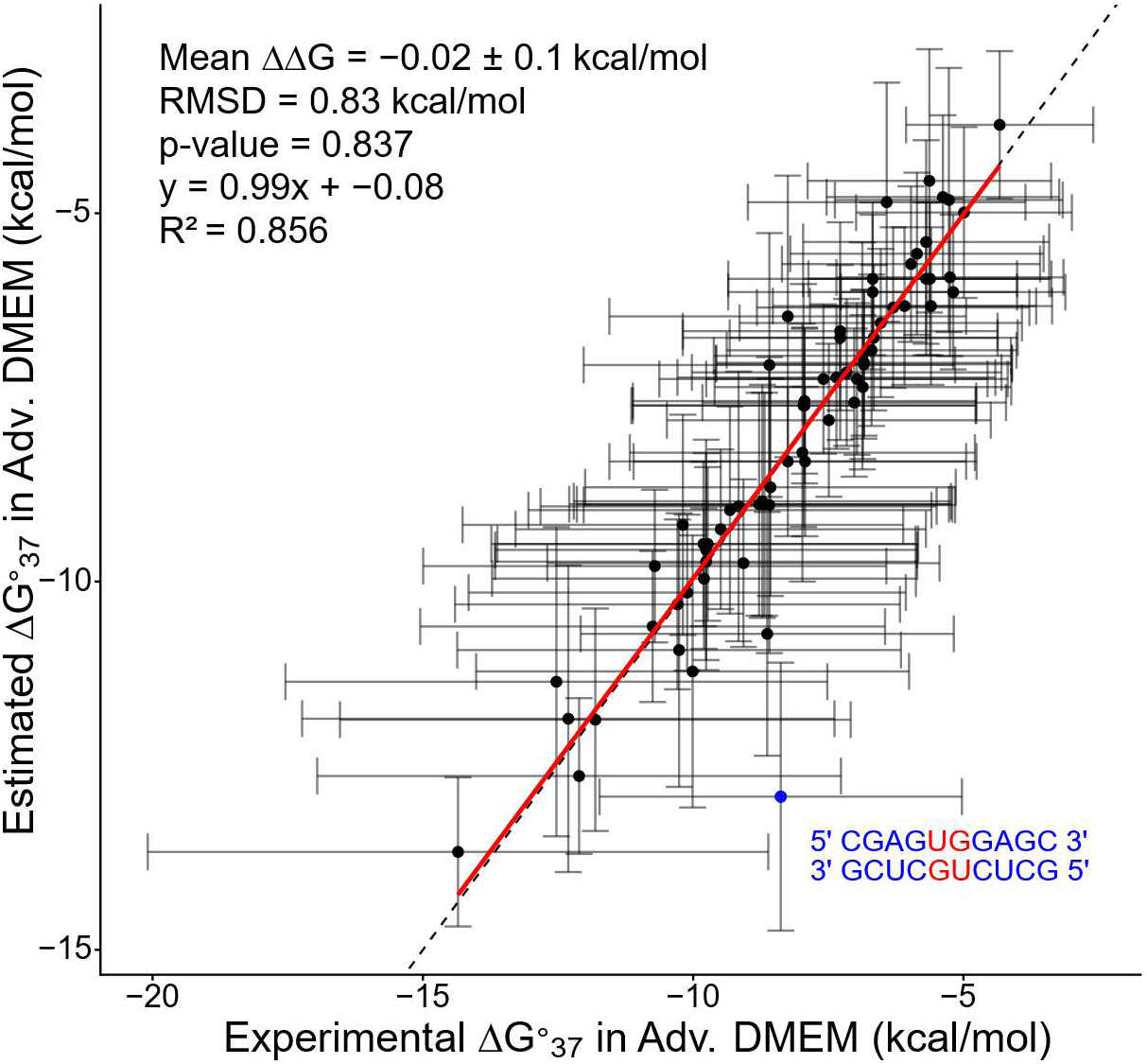
Correlation between experimental and predicted ΔG°_37_ values for RNA helices in Adv. DMEM. Experimental ΔG°_37_ values are plotted against values predicted using the derived Adv. DMEM NN parameters for WCF and G-U stacks. Data closely follow the line of best fit (R^2^ = 0.856), indicating strong overall agreement. The internal 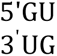 -containing helix (highlighted) is 3^’^UG an outlier, showing substantially lower experimental stability than predicted, likely due to the high uncertainty of tandem 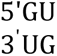 stack parameter in the Adv. DMEM model (Figure 2C and Supplementary Table 4).

### Dangling End Parameters

A set of 32 self-complementary duplexes with dangling ends were studied, each with and without the dangling ends, to determine dangling end parameters in Adv. DMEM (Supplementary Figure 1 and Supplementary Table 5). The sequences were designed to be self-complementary with identical ends to double the stabilizing effect. They include 16 sequences with 5’ dangling ends and 16 sequences with 3’ dangling ends. From these measured folding stabilities, the dangling end free energy contributions were determined using:

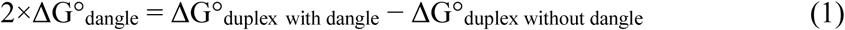

where ΔG°_dangle_ is a parameter that depends on the sequence of the dangling end and its position (on the 5’ or 3’ end of the strand), and the sequence of the adjacent terminal base pair. A table of the Adv. DMEM dangling end parameter results is available (Supplementary Table 6). ΔG°_duplex without dangle_ is the experimental ΔG°_37_ measured for the helix without the dangling end. Parameter uncertainty estimation was carried out using quadrature error propagation [36]. For dangling ends on G-U and U-G pairs in the dataset, we used the Turner 2004 estimates for these parameters.

### Terminal Mismatch Parameters

Terminal mismatch stability parameters depend on the sequence of the mismatch pair and the adjacent closing base pair. A set of 21 self-complementary sequences containing terminal mismatches and their corresponding reference duplexes were melted in Adv. DMEM to obtain their folding free energy changes to calculate the terminal mismatch ΔG°_37_ contributions (Supplementary Figure 2 and Supplementary Table 7). The terminal mismatch stability is determined by:

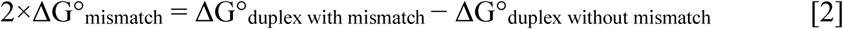

where ΔG°_duplex with mismatch_ is the stability determined by optical melting for the duplex that has the mismatch. ΔG°_duplex without mismatch_ is the experimental stability of the core helix without the terminal mismatch. The factor of two on the left side of the equation is present because the duplexes in this dataset are self-complementary and therefore have the terminal mismatch at both ends of the duplex. The table of terminal mismatch parameters derived in Adv. DMEM is available (Supplementary Table 8). We did not measure all combinations of terminal mismatches and adjacent closing base pairs; therefore, we used the Turner 2004 estimates for missing parameters.

### Hairpin Loops

Hairpin loop stabilities are determined from experiments using:

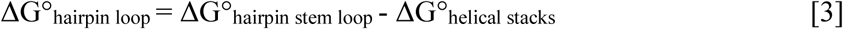

where ΔG°_stem loop_ is the experimentally determined stability of the hairpin with its closing stem and ΔG°_helical stacks_ is the total stability of the helix stacking terms including terminal pair penalties (excluding intermolecular initiation). The set of 17 hairpin sequences and their experimentally determined stability in Adv. DMEM is available (Supplementary Table 9)

The nearest-neighbor model estimates hairpin stability with [25]:

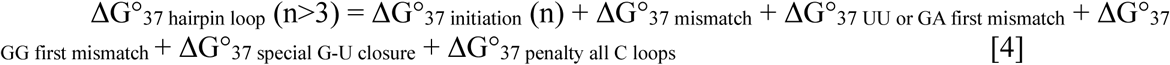

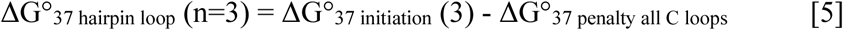

where n is the number of unpaired nucleotides in the loop. ΔG°_37 initiation_ (n) is a sequence-independent term that is the cost for closing the loop of n unpaired nucleotides. ΔG°_37 mismatch_ is the terminal mismatch term for the first mismatch adjacent to the closing base pair. ΔG°_37 UU or GA first mismatch_ and ΔG°_37 GG first mismatch_ are empirical bonuses for UU, GA, or GG first mismatches that were observed in optical melting experiments. Hairpin loops closed by G-U pairs with two preceding G residues receive a bonus, ΔG°_37 special G-U closure_. Loops composed entirely of C residues receive a penalty, ΔG°_37 penalty all C loops_.

Adv. DMEM-specific bonus values for the special first mismatch terms (ΔG°_37 UU or GA first mismatch_ and ΔG°_37 GG first mismatch_) for hairpin loops (n>3) were derived from the experimental hairpin dataset by first calculating the hairpin initiation free energy after accounting for the experimentally measured hairpin stability, the calculated stem free energy, and the corresponding terminal mismatch contribution. The Turner 2004 length-dependent hairpin term was then removed, after correcting the average difference between the calculated hairpin initiation free energy and the Turner 2004 initiation term of non-special hairpins (0.22 ± 0.41 kcal/mol) to isolate the contribution of the special first mismatch bonus. Average bonus values were determined for each special mismatch class and compared with those obtained using 1 M NaCl parameters and the Turner 2004 nearest-neighbor model (Supplementary Table 10).

Sequence-dependent parameterization of these hairpin-specific bonus and penalty terms was also explored. However, only two to three hairpins were available for most mismatch and loop motif classes, and the resulting estimates were inconsistent among sequences. Therefore, although the Adv. DMEM specific average bonus values were derived, they were not incorporated into the final Adv. DMEM parameter set used for RNAstructure benchmarking. Instead, the original Turner 2004 sequence-dependent bonus and penalty terms were retained.

### Internal Loops and Bulge Loops

The stability of a set of 12 sequences containing bulge loops, ranging from 1 to 3 nucleotides, and their reference sequence without the bulge were measured in Adv. DMEM (Supplementary Table 11). A set of 16 symmetric internal loop sequences was also measured in Adv. DMEM alongside their reference sequence (Supplementary Table 12), with the loop size ranging from 1×1, 2×2, and 3×3. The loop stability for bulges and internal loops is determined by:

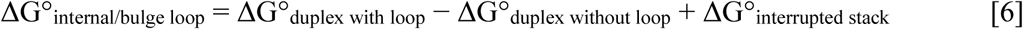

where ΔG°_duplex with loop_ and ΔG°_duplex without loop_ are the stabilities determined by optical melting experiments [46]. ΔG°_interrupted stack_ is the stability of the helical stacking parameter for the stack in the duplex that is interrupted by the loop. The A-U terminal pair penalty is additionally applied to ΔG°_interrupted stack_ per A-U pair in the stack. ΔG°_interrupted stack_ accounts for the fact that the internal loop divides the helix found without the loop into two separate helices. For single-nucleotide bulges, helical stacking is assumed to remain continuous with the bulged nucleotide in solvent, and therefore ΔG°_interrupted stack_ is set to 0 kcal/mol. Bulges with two or more nucleotides, however, are expected to disrupt stacking [25].

### Structure Predictions

Structures were predicted using the RNAstructure software package version 6.6 [26,29]. For our set of sequences with known structures, we used Archive II, which includes a set of 3,960 RNA sequences whose secondary structures have been extensively validated through comparative analysis, including 5S ribosomal RNAs (1283 sequences) [47,48], group I self-splicing introns (98 sequences) [49,50], group II self-splicing introns (11 sequences) [51], large subunit ribosomal RNAs (5 full sequences and 30 domain sequences) [52,53], small subunit ribosomal RNAs (22 full sequences and 88 domain sequences) [54], RNase P RNAs (454 sequences) [55], signal recognition particle (SRP) RNAs (924 sequences) [56], tRNAs (557 sequences) [57], tmRNAs (462 sequences) [58], and telomerase RNAs (37 sequences) [48,59–61].

The *Fold* program in RNAstructure was used with its default settings to predict minimum free energy (MFE) secondary structures for each RNA sequence in Archive II, with the exception of specified folding alphabets: “*rna.”* (1 M NaCl parameter data tables) and “*dmem.”* [23,26–28]. The partition function calculations (generated with program *partition*) estimate base pair probabilities by considering all possible secondary structures weighted by their Boltzmann probabilities [23,26–28]. From these probabilities, the program *MaxExpect* assembles the maximum expected accuracy structure [26,62,63].

Predicted structures were evaluated against known comparative analysis structures using the *scorer* program, which calculates sensitivity (fraction of known base pairs correctly predicted) and positive predictive value (PPV) (fraction of predicted pairs that are correct) [23,26–28].

To further assess the accuracy of RNA secondary structure predictions for sequences in the Archive II benchmarking dataset, we calculated the F1 score for each predicted structure under both the 1 M NaCl and Adv. DMEM nearest-neighbor models. The F1 score was calculated as the harmonic mean of PPV and sensitivity:

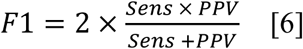

Unlike an arithmetic mean, the harmonic mean penalizes imbalanced performance, such that high PPV with poor sensitivity, or high sensitivity with poor PPV, yields a reduced overall score. The F1 score therefore provides a single-metric assessment of prediction accuracy that balances both correct and comprehensive base-pair recovery and is widely used for evaluating RNA secondary structure prediction methods [63].

The *EDcalculator* program of RNAstructure was used to quantify how well the RNA folds into the known secondary structure as quantified by the normalized ensemble defect (NED). The NED estimates the probability that a nucleotide is misfolded relative to the known structure across the entire thermodynamic ensemble of structures, using base pairing probability estimates [64].

For all accuracy metrics (sensitivity, PPV, F1 score, and NED), values were averaged across sequences within each RNA family. Paired comparisons between the 1 M NaCl and Adv. DMEM conditions were evaluated using one-tailed paired t-tests using the ttest_rel function from SciPy [65], with statistical significance defined as P < 0.05. These tests were used to test the hypothesis that the Adv. DMEM nearest-neighbor model improves RNA secondary structure prediction accuracy relative to the standard 1 M NaCl model.

## Results

### Nearest Neighbor Parameter Fit Procedure

To develop an *in vivo*-like nearest-neighbor model for RNA folding stability in biologically relevant conditions, RNA folding parameters were derived by fitting experimental optical melting data of 193 RNA oligomer model systems containing duplexes and hairpins in Adv. DMEM. We chose the specific sequences following the sensitivity analysis performed for the nearest-neighbor rules in 1 M NaCl, which identified that uncertainty in some features has a larger impact on uncertainty in estimates of base pairing probabilities [36]. We found that, except for internal loops, the functional form of the nearest-neighbor rules determined in 1 M NaCl also works well for Adv. DMEM, when adjustments are made to the parameter values. We fit these values, including dangling ends, terminal mismatches, hairpin loops, and bulge loops, using the established procedures used for the Turner 2004 model.

### RNA Helices are less thermodynamically stable in Adv. DMEM

Agreement between the experimental ΔG°₃₇ values of helices containing WCF and G-U base pairs when measured in Adv. DMEM conditions and those predicted by the Adv. DMEM helical parameters was robust, indicating strong model accuracy for duplex stability (Figure 3, R² = 0.856). The helices included in this correlation analysis were the same duplex dataset used to derive the Adv. DMEM helical parameters. In the correlation plot, predicted values closely tracked experimental measurements across the dataset. One notable outlier was the highlighted internal 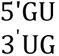 -containing duplex, for which the experimental ΔG°₃₇ was substantially less stable than the corresponding predicted value. This discrepancy may reflect the relatively high uncertainty of the 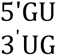 stacking parameter in Adv. DMEM and/or context-dependent effects of tandem G-U pairs, including structural variation or position-specific destabilization not fully captured by the nearest-neighbor model. Similar behavior for tandem G-U motifs has been reported by Zuber et al. [31].

Compared to the Watson-Crick-Franklin (WCF) parameters derived in 1 M NaCl, those obtained in Adv. DMEM showed minimal changes, with an average ΔΔG°₃₇ of 0.05 ± 0.04 kcal/mol across all 10 WCF nearest-neighbor (NN) base pair stacking combinations, the A-U end penalty, and the intermolecular initiation penalty parameters (Figure 2A and B, Supplementary Table 2). This indicates that the change in buffer composition had little impact on these thermodynamic parameters. To ensure a direct comparison and eliminate potential biases from sample size or sequence-specific effects, the ΔG°₃₇ parameters for 1 M NaCl were recalculated using the same 42 helices fitted in Adv. DMEM. The comparison between Adv. DMEM and the refitted 1 M NaCl parameters also showed minimal differences, with an average ΔΔG°₃₇ of 0.02 ± 0.08 kcal/mol across the same WCF parameters (Figure 2A).

To assess global effects on stability, we compared total measured folding stabilities of the helices melted in the two buffers and found that on average, helices melted in Adv. DMEM were about 0.50 ± 0.07 kcal/mol less thermodynamically favorable compared to 1 M NaCl. This difference was statistically significant (Figure 2B: P = 7.68×10^9^, R^2^ = 0.946). These results suggest a destabilizing effect of Ad. DMEM which was not obvious in the nearest-neighbor parameters, potentially due to the large uncertainties involved.

In Adv. DMEM, G-U containing stacks were generally less stable compared to those in 1 M NaCl, with an average ΔΔG°₃₇ of 0.47 ± 0.13 kcal/mol across the 11 G-U NN base pair stacking parameters. The largest destabilizations were observed in tandem G-U stacks (Figure 2C, Supplementary Table 4). When comparing Adv. DMEM to the refitted 1 M NaCl parameters for the same 23 helices, a similar destabilizing effect was observed, with an average ΔΔG°₃₇ of 0.35 ± 0.11 kcal/mol (Figure 2C). However, the relatively large uncertainties in both parameter sets make it challenging to determine the significance of these destabilizing effects.

Direct measurements of helical stability confirmed a statistically significant decrease in G-U-containing helix stability in Adv. DMEM compared to 1 M NaCl. On average, ΔG°₃₇ measurements showed a 0.88 kcal/mol ± 0.12 kcal/mol decrease in stability, consistent with the trend that RNA helices are less stable in Adv. DMEM (Figure 2D: P = 4.51×10^-7^, R² = 0.875).

### Dangling Ends and Terminal Mismatches Have Similar Stability in Adv. DMEM and 1 M Na^+^

Dangling ends are unpaired nucleotides adjacent to helixes and terminal mismatches are non-canonical base pairs adjacent to helices. Both stabilize RNA helices through stacking interactions with adjacent base pairs [66]. In the nearest-neighbor model, the stabilities of these motifs depend on the sequence of the dangling end or terminal mismatch and the sequence of the neighboring base pair. Optical melting experiments in Adv. DMEM were conducted to study 16 dangling end-containing (Supplementary Table 5) and 21 terminal mismatch-containing duplexes (Supplementary Table 7). The *in vivo*-like motif stabilities for dangling ends and terminal mismatches are recorded (Supplementary Tables 6 and 8 respectively).

The derived 5’ and 3’ dangling end parameters in Adv. DMEM closely matched those previously measured in 1 M NaCl, with an average ΔΔG°₃₇ of 0.02 ± 0.05 kcal/mol (Supplementary Figure 1). A two-tailed paired t-test comparing the total ΔG°₃₇ of duplexes containing dangling ends in 1 M NaCl versus Adv. DMEM showed no significant difference, indicating that the stability of dangling ends is consistent across these conditions (P = 0.673). Similarly, the terminal mismatch parameters derived from Adv. DMEM remained largely unchanged compared to 1 M NaCl, with an average ΔΔG°₃₇ of 0.13 ± 0.07 kcal/mol (Supplementary Figure 2). A paired t-test comparing the total ΔG°₃₇ values of duplexes with terminal mismatches in 1M NaCl versus Adv. DMEM also showed no significant difference, suggesting that terminal mismatch stability is unaffected by the change in conditions (P = 0.373)

### Hairpin Loops Trend Less Stable in Adv. DMEM

Hairpin loops are structures formed when a single-stranded RNA folds back on itself, resulting in a helical stem connected by a loop. Using data from 17 hairpins melted in Adv. DMEM, we derived hairpin loop ΔG°_37_ values (Supplementary Table 9). Analyzing the line of best fit for Adv. DMEM versus 1 M NaCl hairpin free energy changes showed that, although the average ΔΔG°_37_ across hairpins was modest, hairpins were systematically less stable in Adv. DMEM, with a fitted intercept of +0.8 kcal/mol, indicating an approximately uniform destabilization across the dataset (Figure 4; P = 0.0291, R² = 0.867). Accordingly, we increased the hairpin initiation terms from the 1 M NaCl model by 0.8 kcal/mol for the Adv. DMEM model.

**Figure 4.**
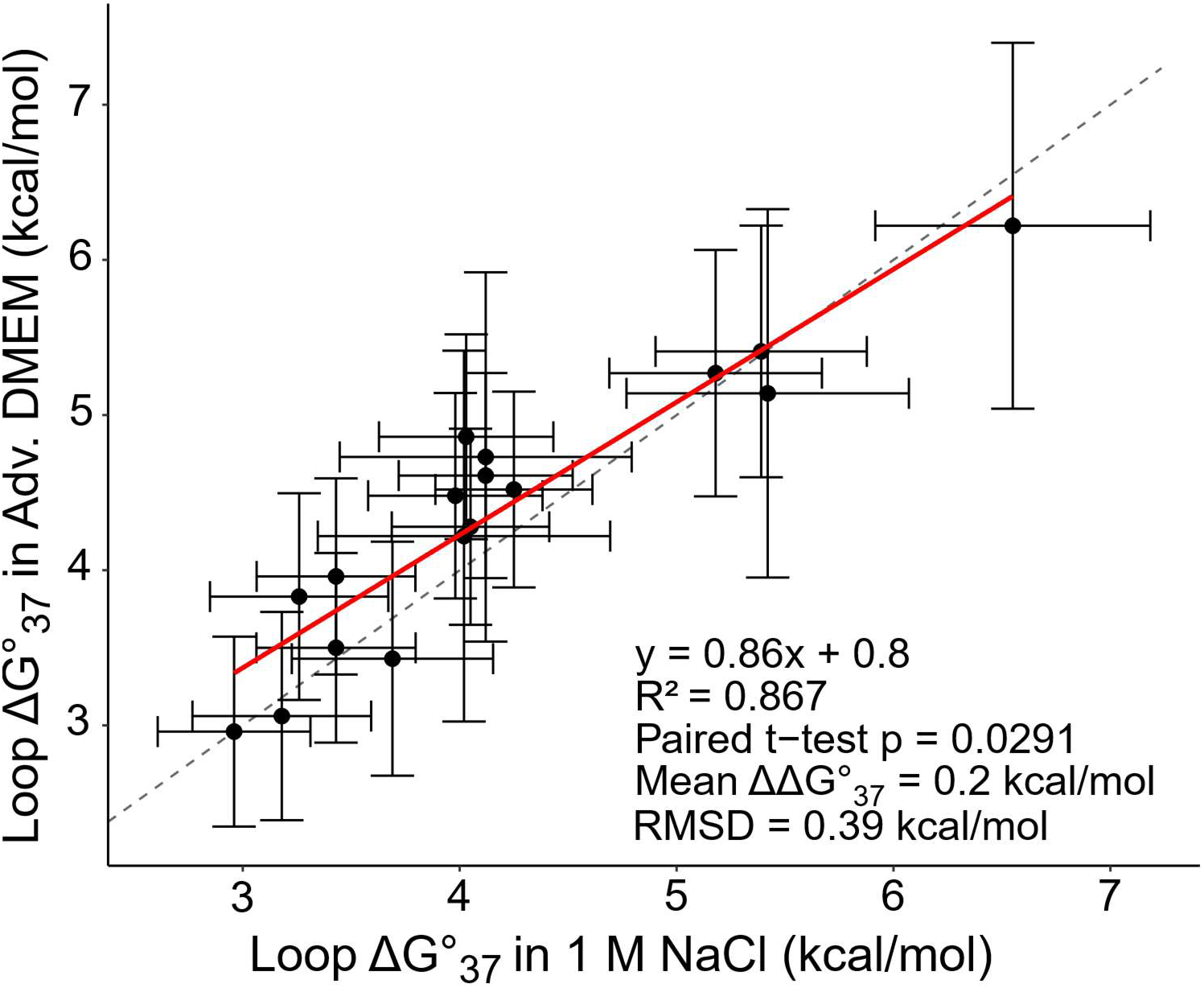
ΔG°_37_ of hairpin loops measured via UV optical melting in Adv. DMEM versus 1M NaCl. Points and error bars represent the loop folding ΔG°_37_ and uncertainty of 17 hairpin loops in Adv. DMEM versus 1 M NaCl [31]. Hairpin loop ΔG°_37_ is determined by subtracting the helix stem ΔG°_37_ from the hairpin ΔG°_37_. For loops of four or more nucleotides, terminal mismatch contributions are also subtracted. The R^2^ represents the coefficient of determination of the linear model presented by the figure and represented graphically with the dashed line. The solid line represents the diagonal. The two-tailed paired t-test between the folding stabilities of the helices measured in the two conditions was performed using the t.test function in R.

### Bulge Loops of One Nucleotide Are Less Stable in Adv. DMEM

Bulge loops are unpaired nucleotides that interrupt a helix. The thermodynamic cost of forming bulges, estimated by the loop initiation parameters, is length specific. For single-nucleotide bulges, helical stacking is assumed to remain continuous, whereas bulges with two or more nucleotides disrupt stacking [25]. Optical melting experiments in Adv. DMEM were conducted with 12 bulge loop models with loop sizes ranging from one to three unpaired nucleotides (Supplementary Table 11). Bulge loop initiation values for two-and three-nucleotide loops were consistent with the 1 M NaCl initiation values, while single-nucleotide bulges were 1.59 ± 0.41 kcal/mol less favorable in Adv. DMEM (Figure 5).

**Figure 5.**
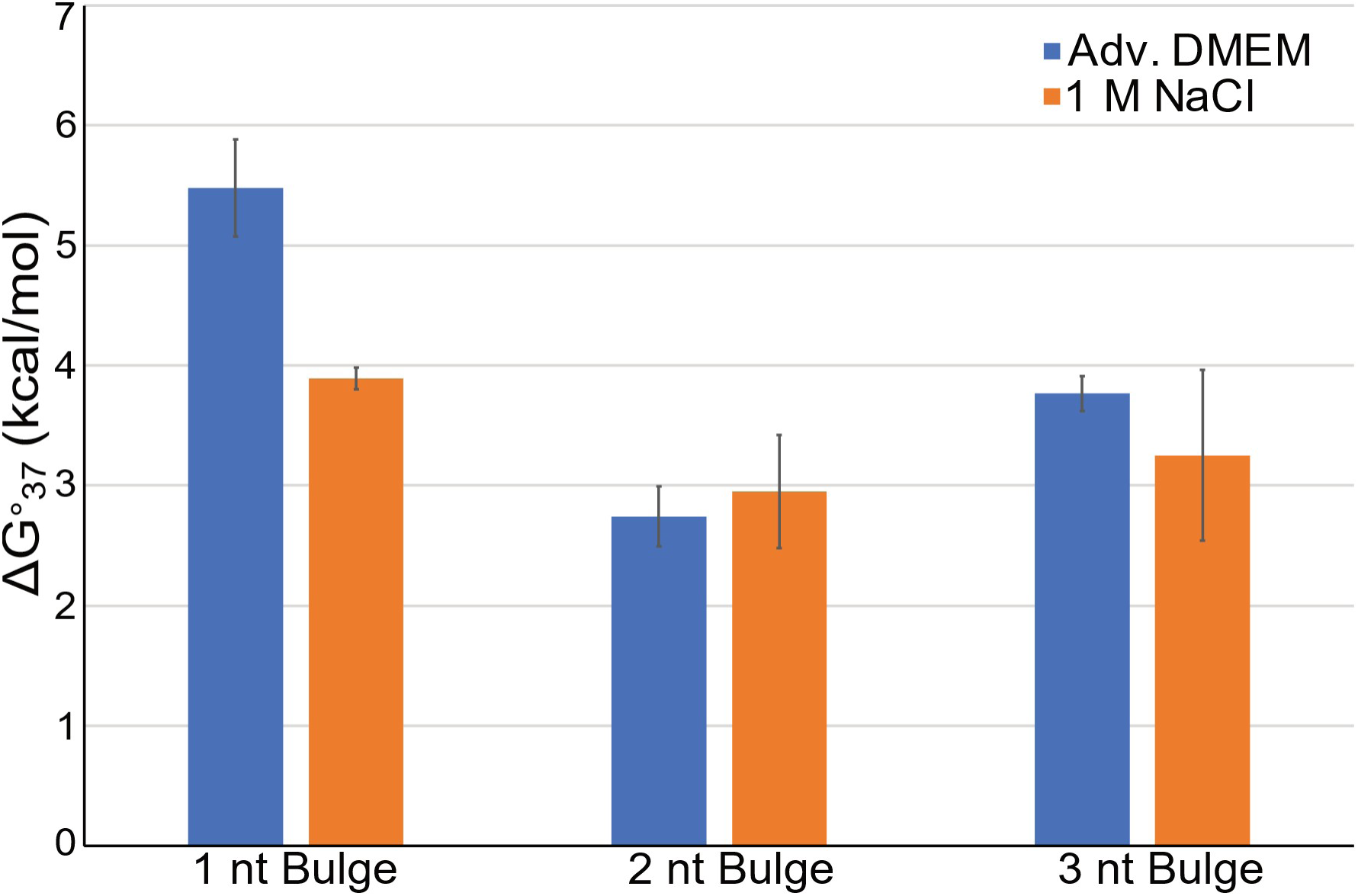
Comparison of bulge loop parameters in Adv. DMEM versus 1 M NaCl. ΔG°_37_ values for bulge loops of one to three nucleotides were derived by subtracting the ΔG°_37_ of the core helix (or helices, for loops >1 nucleotide) from the measured duplex in Adv. DMEM (blue) and compared to established bulge loop parameters in 1 M NaCl (orange) [31]. Error bars for Adv. DMEM indicate uncertainty estimated by standard error of the mean. A paired t-test comparing total ΔG°₃₇ values for the bulge-containing duplexes across all three lengths in the two buffers showed no significant difference (P = 0.054).

### Internal Loops Display Idiosyncratic Behavior in Adv. DMEM

Internal loops consist of unpaired nucleotides on both strands that disrupt the double helix. The loop stabilities were determined for 16 symmetric internal loops (1×1, 2×2, and 3×3 loops) in Adv. DMEM (Supplementary Table 12). Except for pyrimidine-only mismatches, which exhibited a stabilizing contribution, internal loops in Adv. DMEM were generally destabilized compared to 1 M NaCl, with the average ΔΔG°₃₇ of -0.93 ± 0.54 kcal/mol between duplexes with internal loops melted in Adv. DMEM vs 1 M NaCl (Figure 6: R² = 0.034). However, 2×2 and 3×3 all-uracil loops remained unchanged in stability.

**Figure 6.**
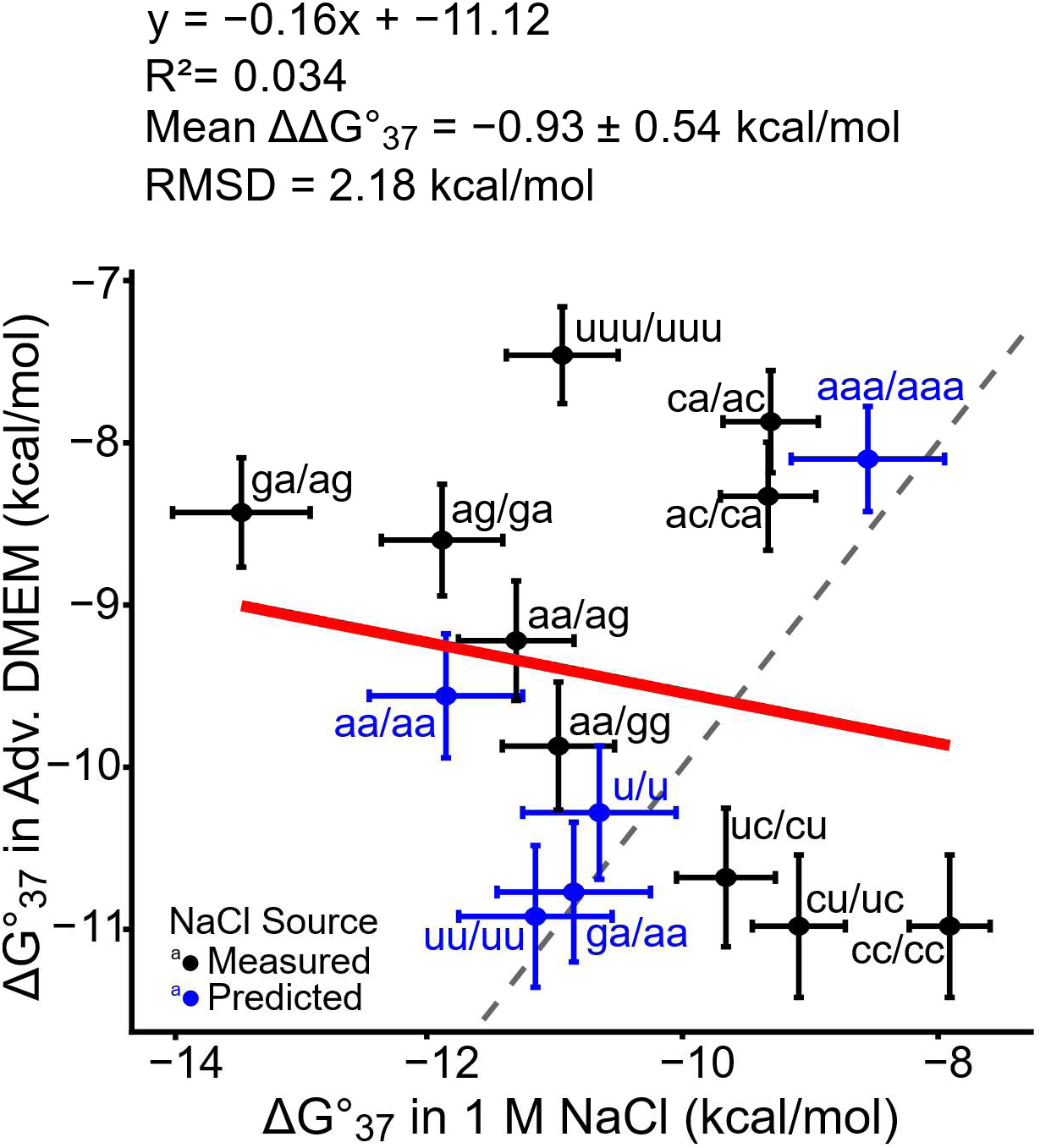
ΔG°_37_ of internal loops measured via UV optical melting in Adv. DMEM versus 1M NaCl. Total duplex ΔG°_37_ for 16 internal loops in Adv. DMEM as a function of ΔG°37 in 1 M NaCl [31]. The R^2^ represents the coefficient of determination of the linear fit (red line). The dashed line represents the diagonal.

High uncertainties in the derived internal loop parameters and poor correlation with 1 M NaCl values indicate that stability trends in physiological-like medium are largely sequence-and context-dependent. As a result, a robust NN model for internal loops could not yet be established. For the current time, the Adv. DMEM parameters still use the internal loop parameters for 1 M Na^+^, which were derived using 304 optical melting experiments [25].

### Multibranch Loop Parameters

We chose to not perform experiments at this time on multibranch loops. In the 1 M NaCl model, multibranch-loop free energies are described using a simple linear model based on the number of branches and unpaired nucleotides, with parameters fit to optical melting data [67,68]. Previous work comparing alternative multibranch-loop models found that this conventional linear model performed as well as, or better than, more complex polymer-physics-based models for RNA secondary structure prediction [69]. Subsequent parameter optimization by maximizing the accuracy of secondary structure prediction further showed that the established linear-model parameters are near optimal [70]. Thus, given that the simple linear model for multibranch loops works well for structure prediction, we retained the established 1 M NaCl multibranch-loop parameters.

### Secondary Structure Prediction

For software implementation and benchmarking, we modified the RNAstructure parameter libraries and stored the resulting model as a new folding alphabet, “*dmem*.” The “*dmem.”* folding alphabet uses the derived Adv. DMEM parameters for canonical helical base stacks and secondary structure loops. Then, to test the biological relevance of the new Adv. DMEM nearest-neighbor model, we used them for secondary structure prediction for the ten families of RNA sequences and structures in the Archive II dataset [61].

The predictive performance of the Adv. DMEM model was evaluated using two approaches. First, we predicted minimum free energy (MFE) structures [23,26–28]. Second, we predicted maximum expected accuracy structures (MEA) [23,26–28]. Both approaches were applied using folding parameters derived from the 1 M NaCl model and the *in vivo*-like Adv. DMEM model. The predicted structures were then compared to the corresponding accepted structures determined by comparative sequence analysis, using three metrics: (i) Sensitivity, which measures the fraction of known pairs correctly predicted, (ii) Positive predictive value (PPV), which represents the fraction of predicted pairs that are in the known structure, and (iii) F1 scores which is the harmonic mean of PPV and sensitivity.

Predictions of MFE and MEA structures using Adv. DMEM folding parameters showed no substantial improvement in overall accuracy across RNA families in Archive II. On average, the sensitivity of structures predicted with the Adv. DMEM model was lower than that of structures predicted with the 1 M NaCl model (Supplementary Figure 3A and 4A; Supplementary Tables 13 and 14). In contrast, PPV showed a modest increase when using *in vivo*–like conditions (Supplementary Figure 3B and 4B; Supplementary Tables 13 and 14). Consistent with these opposing trends, F1 scores, which integrate sensitivity and PPV into a single accuracy metric, were generally similar between models for both MFE and MEA predictions, with only modest improvements observed in selected RNA families (Supplementary Figure 3C and 4C; Supplementary Tables 13 and 14). Together, these results suggest that, although the Adv. DMEM model may slightly reduce the prediction of false-positive base pairs, it does not improve the sensitivity of predictions and therefore requires further refinement before widespread application.

tRNA sequences were a notable exception to this overall trend. For tRNAs, the Adv. DMEM model significantly improved PPV for MFE structures, with an approximately 3% increase relative to the 1 M NaCl model (Supplementary Figure 3B; P = 0.008; Supplementary Table 13). This improvement was also reflected in the F1 score, which increased by approximately 1.77% under the *in vivo*-like Adv. DMEM model (Supplementary Figure 3C; P = 0.011; Supplementary Table 13). The higher F1 score indicates improved overall prediction accuracy, reflecting a better balance between recovery of true base pairs and reduction of false-positive predictions for tRNA structures.

### Adv. DMEM Model Enhances Desired Secondary Structure Folding

To evaluate how effectively the Adv. DMEM nearest-neighbor model improves RNA folding toward the correct secondary structure in the thermodynamic ensemble, we estimated the normalized ensemble defect (NED) score, which is the average probability that a nucleotide is misfolded relative to the reference structure across the entire thermodynamic ensemble of structures [64]. A lower normalized NED score indicates that the native structure, such as the cloverleaf for tRNA, is more strongly favored in the ensemble.

In this analysis, the known structure was defined as the base-pairing obtained through comparative sequence analysis. Results showed that the average NED scores for most RNA families in the Archive II set were modestly improved with the Adv. DMEM model. However, 16S (using domain sequences), group 2 self-splicing introns, telomerase RNA, tmRNA, and tRNA exhibited significantly lower NED scores under the Adv. DMEM model compared to the 1 M NaCl model, indicating improved folding accuracy for these RNA types (Figure 7; P = 0.0158, P = 0.0014, P = 0.0139, P = 0.0051, P = 2.21×10^-33^, respectively).

**Figure 7.**
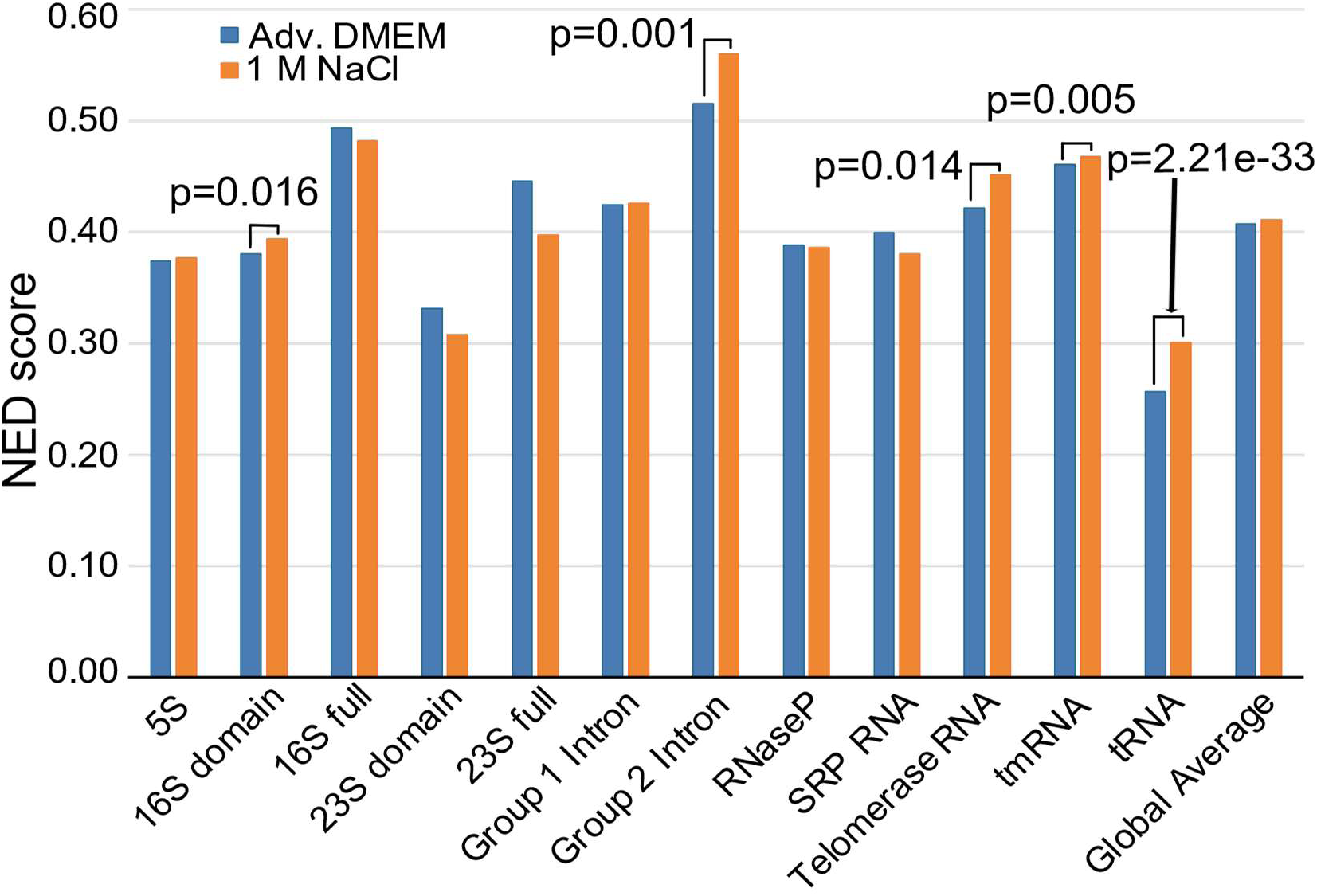
Comparison of average normalized ensemble defect (NED) scores across 10 RNA families under the Adv. DMEM nearest-neighbor model vs 1 M NaCl nearest-neighbor model. Significant improvements in NED scores under the Adv. DMEM model are labeled and were determined using one-sided paired t-tests.

## Discussion

### RNA Helices are Less Stable *in vivo*

RNA helices demonstrate decreased thermodynamic stability with *in vivo*-like conditions compared to measurements conducted in 1 M NaCl. In Adv. DMEM. We observed that helices of WCF pairs and helices with G-U including pairs, on average, were 0.5 kcal/mol and 0.9 kcal/mol less stable, respectively, than in 1 M NaCl (Figure 2B and 2D). These findings align with the observations recently reported by Sieg *et al.* (2023), which documented reduced RNA helix stability in Eco80, an artificial *in vivo*-like medium designed to mimic *E. coli* cellular conditions [33]. Because Eco80 is opaque at the optical melting wavelengths (260 and 280 nm), fluorescence-detected binding isotherms (FDBI) were employed to assess RNA helix stability in Eco80. These measurements require covalently attached fluorophores and quenchers.

A comparison of WCF nearest-neighbor parameters derived from Adv. DMEM and Eco80 reveals notable differences in stability for specific nearest-neighbor WCF pairs (Supplementary Figure 5). For instance, FDBI-derived parameters indicate weaker stability for 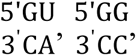 and 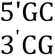 nearest-neighbors compared to those obtained by UV optical melting. These discrepancies underscore methodological differences between FDBI and UV optical melting experiments. FDBI is complicated by the potential effects of fluorophore and quencher labels on RNA folding and interactions with salts and biomolecules at helix termini. Therefore, stabilizing or destabilizing contributions from label stacking must be accounted for when deriving nearest-neighbor parameters under this method. However, Eco80 is a better mimic of intracellular conditions than Adv. DMEM, which is meant to mimic extracellular conditions.

Adv. DMEM optical melting experiments offer a promising avenue for generating extensive datasets to better understand RNA folding behavior in an *in vivo*-like environment. While Eco80 provides a more intracellular-like medium, its methodological limitations may restrict data collection depth. There is an opportunity to integrate findings from diverse experimental approaches to elucidate RNA folding stability *in vivo*. Together, this current evidence suggests that RNA helices in cellular-like environments are less stable, with G-U pairs and loops being particularly sensitive to these in vivo conditions. Collaborative efforts will be essential to integrate the complex interplay of factors influencing RNA stability *in vivo*.

### tRNA Folding Accuracy and the Need for Internal Loop Refinement

Benchmarking results demonstrate that the Adv. DMEM model significantly improves PPV and F1 scores for MFE structure predictions for tRNA (Supplementary Figure 4B and 4C) as well as significantly improves the NED score for several RNA families (specifically 16S rRNA (domain sequences), group 2 introns, telomerase RNA, tmRNA and tRNA) (Figure 7). These findings highlight the potential of the Adv. DMEM model to better replicate RNA folding behavior in cellular-like environments, particularly for RNA families. However, despite these advancements, sensitivity metrics for structure predictions are lower on average with the Adv. DMEM model compared to the 1 M NaCl Turner 2004 model (Supplementary Figures. 4A and 5A). This suggests that while the Adv. DMEM thermodynamic model enhances the accuracy of correctly predicted base pairs, it fails to capture a subset of true base pairs, indicating the need for further refinement.

Among RNA families, tRNAs exhibit the most pronounced improvement in predictive accuracy. For tRNAs, F1 scores increased by almost 2% for MFE structures (Supplementary Figure 4C), while NED scores showed a dramatic reduction from 0.301 to 0.257 (P = 2.21×10^-33^) (Figure 7). These results demonstrate enhanced reliability and precision in secondary structure predictions under *in vivo*-like conditions. A representative example using an isoleucine tRNA in *Mycoplasma capricolum*, showing the significant improvement in folding accuracy under the Adv. DMEM model, is presented in Figure 8. The MEA structure predicted using the 1 M NaCl model deviated substantially from the canonical cloverleaf architecture and contained multiple incorrect base pairs (Figure 8A and Figure 8B), resulting in lower sensitivity (52.38%), lower PPV (45.83%), and a higher NED score (0.362) (Figure 8D). In contrast, the Adv. DMEM model accurately recovered the canonical cloverleaf fold with nearly complete prediction of accepted base pairs (Figure 8A and 8C), yielding markedly improved sensitivity (95.24%), PPV (100%), and NED (0.050) (Figure 8D). Base-pair probability annotations further indicated stronger thermodynamic support for native structure formation under the Adv. DMEM model. Collectively, these results demonstrate substantially improved reliability and precision of tRNA secondary structure prediction under *in vivo*-like conditions. A second example of a serine tRNA from *Candida cylindracea* is shown in Supplementary Figure 6.

**Figure 8.**
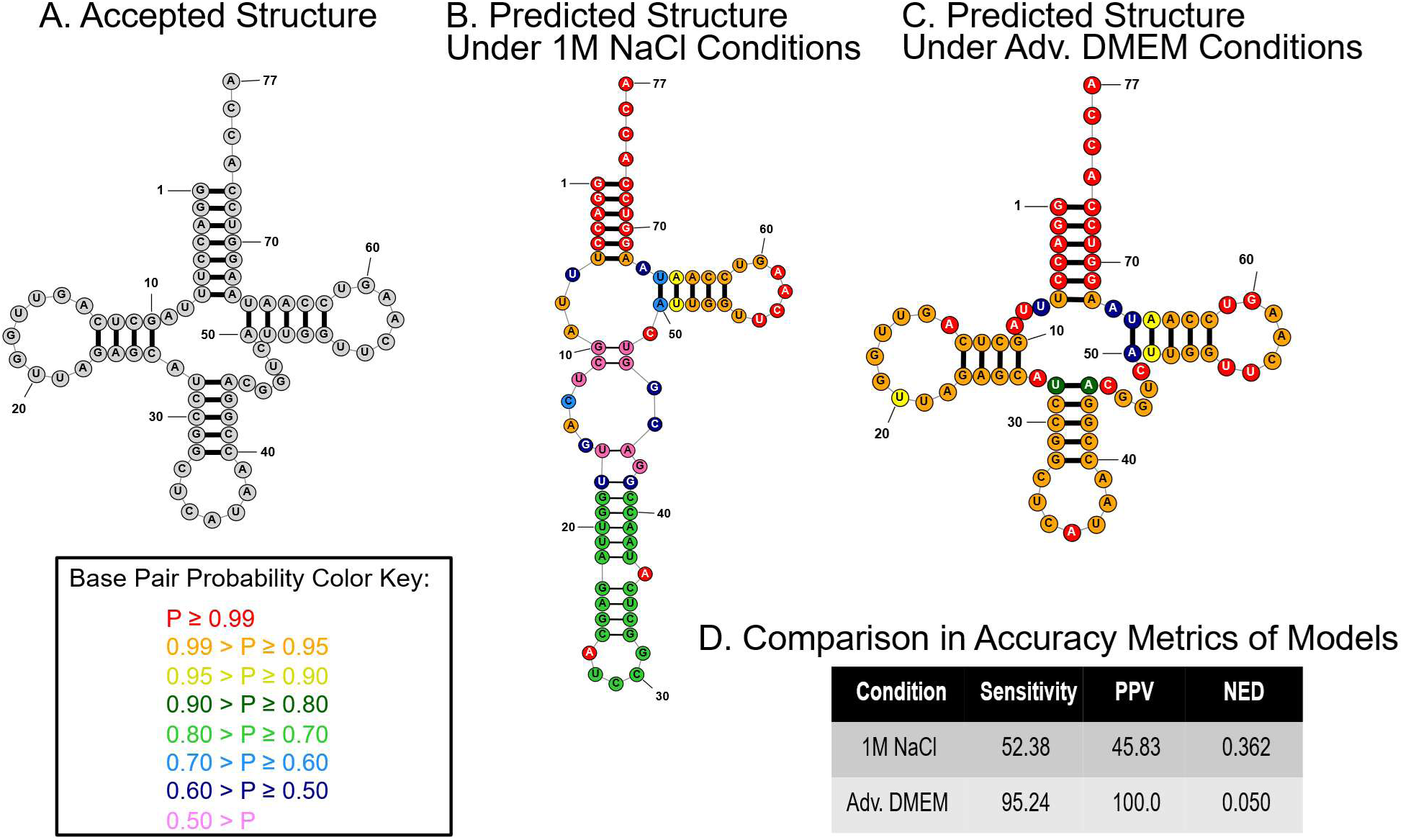
Comparison of MEA structure predictions for a representative tRNA sequence, tdbR00000153-Mycoplasma_capricolum-2095-Ile-AU, under Adv. DMEM and 1 M NaCl nearest-neighbor models. **(A)** Accepted tRNA secondary structure based on the comparative-analysis reference structure. **(B)** MEA predicted secondary structure using the 1 M NaCl nearest-neighbor model. **(C)** MEA predicted secondary structure using the Adv. DMEM model. **(D)** Table comparing sensitivity, positive predictive value (PPV), and normalized ensemble defect (NED) for the two model predictions for this tRNA sequence. All structures were generated with RNAstructure. Base pairs matching the accepted comparative structure are shown with thick lines, whereas incorrect predicted base pairs are shown with thin lines. Base pairs are additionally color-coded according to their computed pairing probabilities.

### Need for Internal Loop Measurements

Interestingly, tRNAs typically lack internal loops, which are not yet modeled in Adv. DMEM. This observation suggests that *in vivo-*like conditions may provide additional precision in secondary structure predictions that the current *in vitro* model lacks. However, this effect may be obscured in RNA families other than tRNAs due to the continued use of 1 M NaCl parameters for internal loops in the Adv. DMEM model. Future studies should focus on determining and modeling internal loop stabilities in Adv. DMEM. We hypothesize that a refined model for internal loops in Adv. DMEM would improve predictions for RNA families with diverse structural features.

### Limitations for Adv. DMEM Model

Although the Adv. DMEM model was designed to better approximate physiological ionic conditions than traditional high-salt buffers, it remains an incomplete representation of intracellular RNA folding environments [71]. First, Adv. DMEM more closely resembles serum or extracellular fluid than the cytosol, as its composition is based on cell-culture medium containing physiologically relevant inorganic ions, nutrients, and metabolites, but lacking the tightly regulated compartment-specific chemistry of living cells (Table 1). The D2902 Adv. DMEM formulation used in this study closely approximates physiological extracellular concentrations of major ions, including Na⁺ (∼141–155 mM), K⁺ (5.4 mM), and Mg²⁺ (0.81 mM) [72]. Glucose concentration (5.55 mM) is also comparable to that of healthy human serum (∼5.0 mM) [72]. In contrast, most amino acids are present at higher concentrations compared to physiological levels, typically 2–13-fold higher than serum, while glutamine (4.0 mM) and pyruvate (1.0 mM) are enriched approximately 8-fold and 29-fold, respectively [72]. Adv. DMEM reproduces the ionic environment of serum reasonably well but provides substantially greater nutrient availability than is present in serum.

Second, both Adv. DMEM and 1 M NaCl conditions are relatively dilute solutions that do not reproduce the extensive macromolecular crowding present *in vivo*. It has been suggested that the total concentration of macromolecules reaches 400 mg/ml and that 40% of the intracellular space is occupied by biomacromolecules [73]. RNA folding thermodynamics has been studied under simulated cellular crowding, typically using polyethylene glycols (PEGs) to mimic the dense intracellular environment [34,35,74]. A significant finding from those studies is that molecular crowders, particularly low-molecular-mass species like PEG200, often destabilize RNA secondary structures, contrary to hierarchical models where strong secondary structures are assumed to form first [75]. This behavior has led to the derivation of specialized base pairing nearest-neighbor parameters to account for environmental effects not captured by the standard 1 M NaCl model. A study in 2019 from Adams & Znosko, fit helical nearest-neighbor parameters for RNA in 1 M NaCl with 20 wt% PEG200. They found that duplexes are destabilized by an average of 1.02 kcal/mol compared to standard buffer conditions [35]. Ghosh *et al.* further developed parameters for physiological 100 mM NaCl with and without 40 wt% PEG200; creating a “universal” framework that separates energetic contributions into bulk interactions, cation effects, excluded volume and water activity [34]. However, there is evidence to suggest that PEG200 and other lower molecular mass PEGs are not an accurate cellular crowding agent and do not mimic physiological crowding conditions because they destabilize RNA structures and disrupt ligand binding through strong attractive interactions, which differ from the stabilized effects observed in cells [74].

Comparing Adv. DMEM parameters to those derived under crowding conditions revealed that the relative ordering of stack stabilities is largely identical across environments (Supplementary Figure 3). Notably, the Adv. DMEM parameters most closely resembled those determined in 100 mM NaCl without crowders from the Ghosh *et al.* study, whereas the inclusion of PEG200, particularly at 40 wt%, increased helix initiation and terminal penalties and altered the magnitude of multiple stacking interactions (Supplementary Figure 3). These limitations indicate that, while Adv. DMEM provides a useful intermediate model between standard *in vitro* buffers and biological environments, future parameterization efforts incorporating molecular crowding and intracellular-like compositions might be necessary to more fully capture *in vivo* RNA thermodynamics.

### Applications

Nearest neighbor folding parameters are the thermodynamic foundation for not only RNAstructure, but also many other RNA folding packages [26,29,76–79]. By grounding these parameters in a more physiologically relevant environment, we can improve the biological relevance of RNA structure predictions. This Adv. DMEM model derived from this study establishes a foundation for studying RNA folding under *in vivo*–like conditions, which has previously been challenging.

An example of an application of extracellular folding conditions is glycoRNAs, which are small noncoding RNAs modified with N-glycans and displayed on the surface of living mammalian [80]. The folding stability of these RNAs is the extracellular ionic environment, which is a direct application of the Adv. DMEM parameters. Recent advances in glycoRNA detection further highlight the potential utility of the Adv. DMEM model. Gong *et al.* developed GLINT; a highly sensitive imaging platform that revealed glycoRNA localization in lipid raft domains, SNARE-mediated transportation to the cell surface and distinct glycoRNA signatures across breast cancer subtypes [81]. Because the RNA component is a major source of glycoRNA heterogeneity, more accurate secondary structure predictions under extracellular-like conditions could improve our understanding of glycoRNA function and transport. Incorporating Adv. DMEM-derived thermodynamic parameters into RNA folding algorithms may also facilitate the discovery of novel glycoRNA structural motifs and improve probe design for technologies such as GLINT by identifying accessible RNA regions for hybridization. Ultimately, these advances could aid the identification of disease-associated glycoRNA signatures and enhance future diagnostic applications.

### Adv. DMEM Parameter Dissemination

The Adv. DMEM nearest-neighbor parameter tables are available in plain-text format as part of the RNAstructure software package (https://rna.urmc.rochester.edu/RNAstructure.html), enabling their use in secondary structure prediction [26,29]. As with the existing 1 M Na+ Turner 2004 parameters, the new Adv. DMEM parameters are compatible with all RNAstructure command line and tools. These *in vivo*-like folding parameters can be specified in RNAstructure using the --alphabet parameter which can switch the canonical RNA alphabet to the alternative Adv. DMEM when specified (invoked with --alphabet dmem).

## Acknowledgements

This work was supported by National Institutes of Health grant R35GM145283 to DHM. OH was partially supported by National Institute of Health training grant T32GM135134. Research was also supported by National Science Center of Poland grants 2022/45/B/ST4/03586 to R.K. and 2021/41/B/NZ1/03819 and 2020/39/B/NZ1/03054 to E.K.

