## Supplemental Information for "Nearest Neighbor Parameters for Estimating RNA Folding Stability with *In Vivo*-like Conditions"

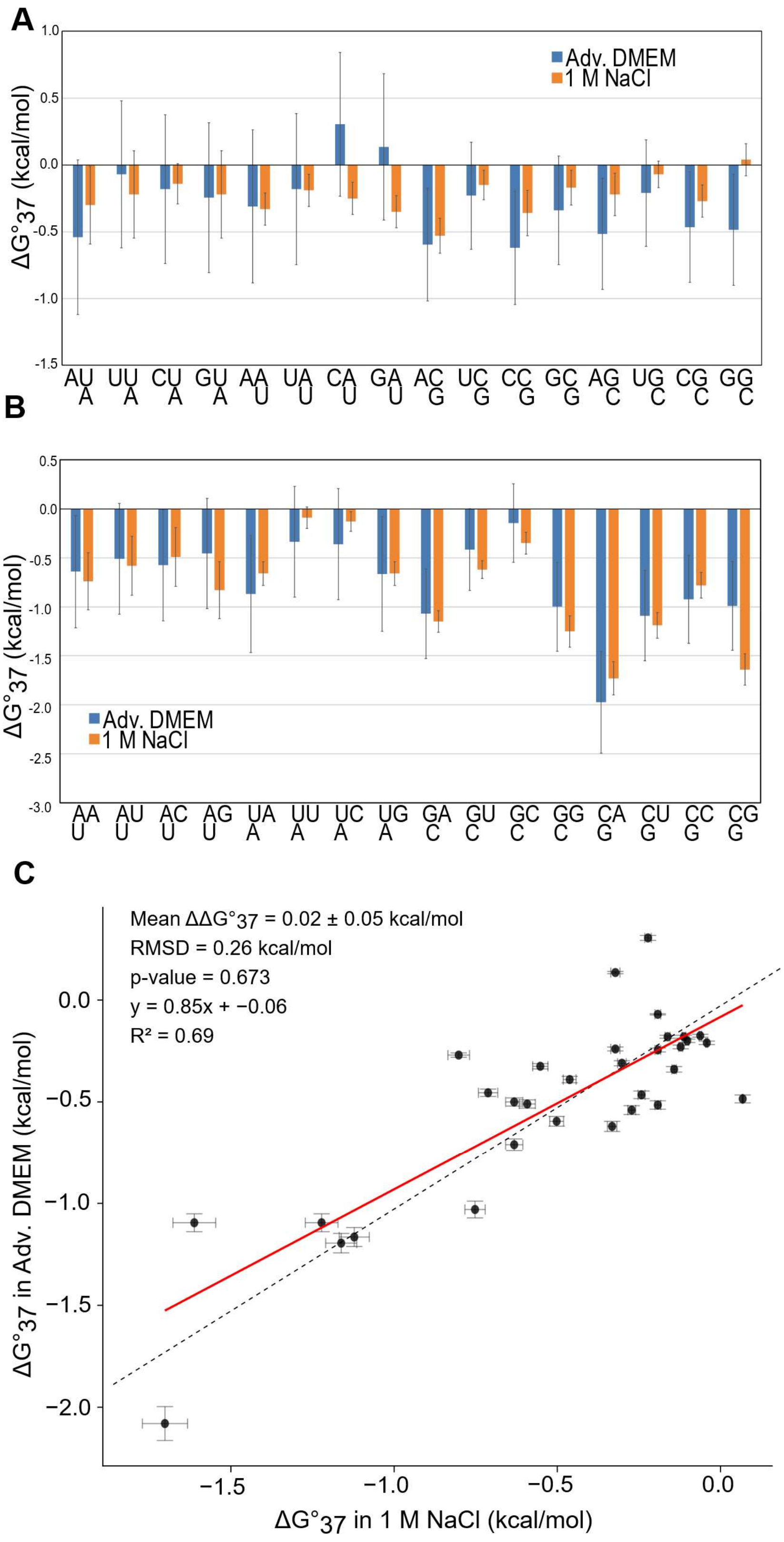

**Supplementary Figure 1. Comparison of dangling end parameters in Adv. DMEM versus 1M NaCl. (A)** $\Delta G^\circ_{37}$ values for 5' dangling ends in Adv. DMEM (blue) compared to established 5'

dangle end parameters in 1M NaCl (orange) [1]. Error bars indicate parameter uncertainty estimated by quadrature error propagation. (**B)** $\Delta G°_{37}$ values for 3' dangling ends in Adv. DMEM (blue) compared to established 3' dangle end parameters in 1M NaCl (orange) [1]. Error bars indicate parameter uncertainty estimated by quadrature error propagation. (**C)** Comparison of total $\Delta G°_{37}$ of duplexes with dangling ends measured in Adv. DMEM versus 1 M NaCl via UV optical melting experiments. Points represent the $\Delta G°_{37}$ and parameter uncertainty of 32 duplexes with dangling ends, 16 duplexes with 5' dangling ends and 16 duplexes with 3' dangling ends, derived in Adv. DMEM versus 1M NaCl [1,2]. The $R^2$ represents the coefficient of determination of the linear model presented by the figure and represented graphically with the red line. The dashed line represents y = x. A paired t-test comparing total $\Delta G°_{37}$ values for dangling end duplexes in the two buffers showed no significant difference. Error bars represent experimental uncertainty.

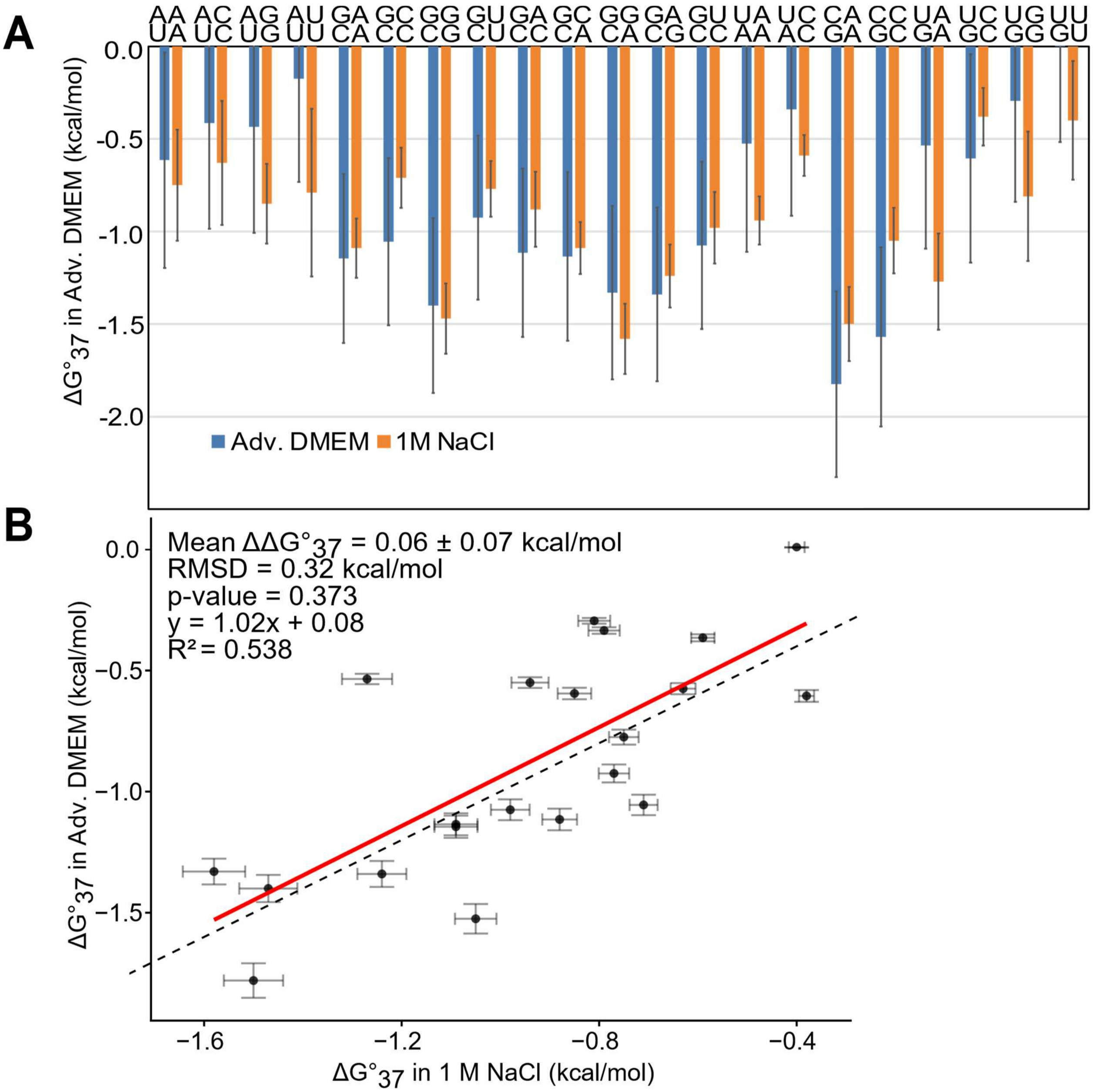

**Supplementary Figure 2. Comparison of terminal mismatch parameters in Adv. DMEM vs 1M NaCl. (A)** $\Delta G°_{37}$ values for terminal mismatches in Adv. DMEM (blue) compared to established terminal mismatch parameters in 1M NaCl (orange) [2]. Error bars indicate parameter uncertainty estimated by quadrature error propagation. (**B)** Comparison of total $\Delta G°_{37}$ of duplexes with terminal mismatches measured in Adv. DMEM versus 1 M NaCl via UV optical melting experiments. Points represent the $\Delta G°_{37}$ and parameter uncertainty of 21 self-complementary duplexes with terminal mismatches, derived in Adv. DMEM versus 1M NaCl [2].The $R^2$ represents the coefficient of determination of the linear model presented by the figure and represented graphically with the red line. The dashed line represents the diagonal. A paired t-test comparing total $\Delta G°_{37}$ values for duplexes with terminal mismatches in the two buffers showed no significant difference. Error bars represent experimental uncertainty.

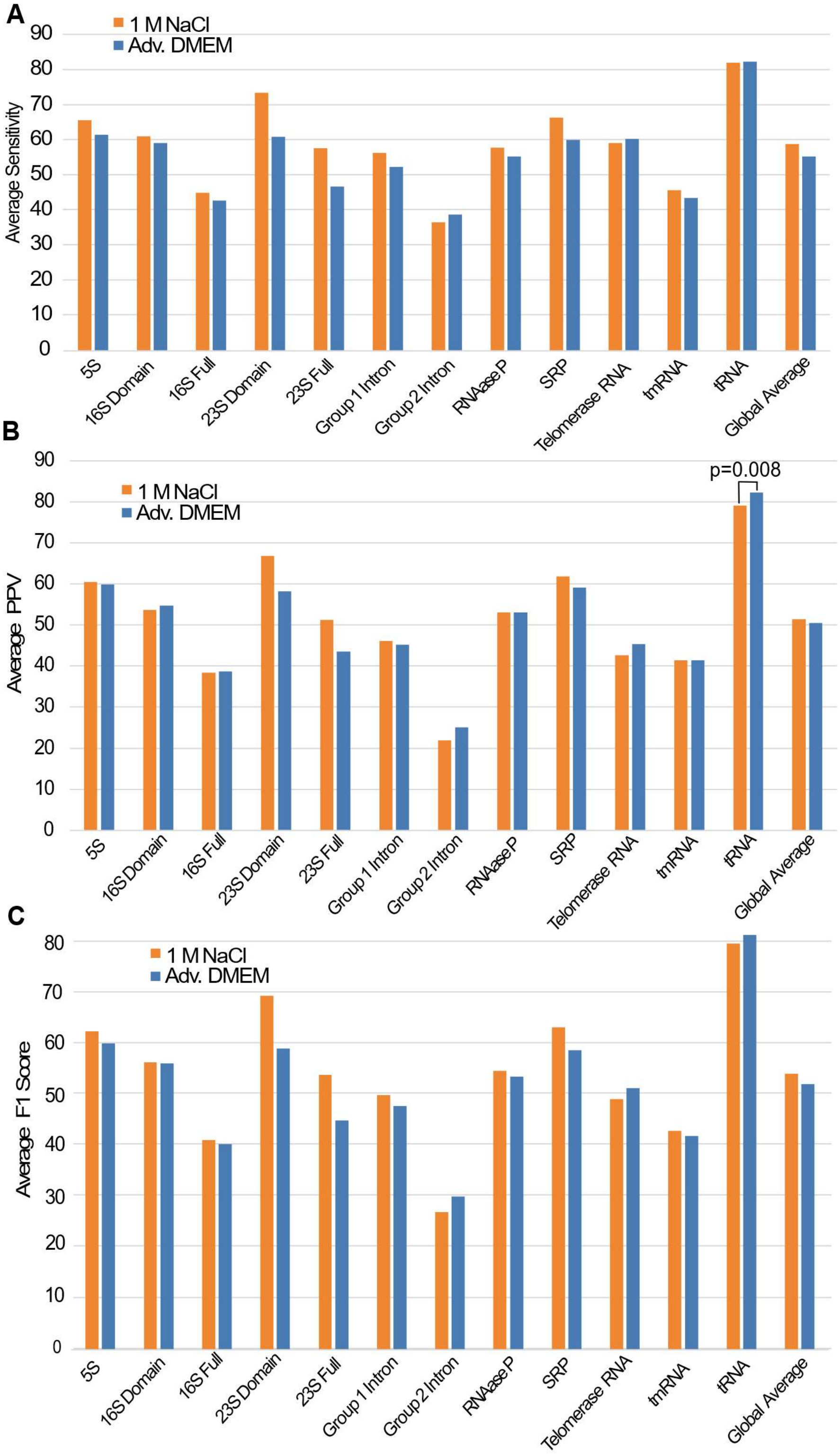

**Supplementary Figure 3. Comparison of MFE secondary structure prediction accuracy using standard 1 M NaCl and Adv. DMEM nearest neighbor parameters. (A)** Average sensitivity scores of minimum free energy (MFE) RNA secondary structure predictions across ten ncRNA families using standard 1 M NaCl nearest neighbor parameters (orange) and Adv. DMEM-derived parameters (blue). For 16S and 23S rRNAs, predictions were evaluated separately for full-

sequence structure prediction and prediction of individual folding domain structures [3,4]. One-sided paired t-tests were performed to assess whether structure predictions improved using Adv. DMEM parameters; significant ($P<0.05$) P-values are indicated.
**(B)** Average positive predictive value (PPV) scores of MFE RNA secondary structure predictions across the same ncRNA families and parameter sets described in **(A)**. One-sided paired t-tests were performed to assess whether structure predictions improved using Adv. DMEM parameters; significant ($P<0.05$) P -values are indicated.
**(C)** Average F1 scores of MFE RNA secondary structure predictions across the same ncRNA families and parameter sets described in **(A)**. One-sided paired t-tests were performed to assess whether structure predictions improved using Adv. DMEM parameters; significant ($P<0.05$) P -values are indicated.

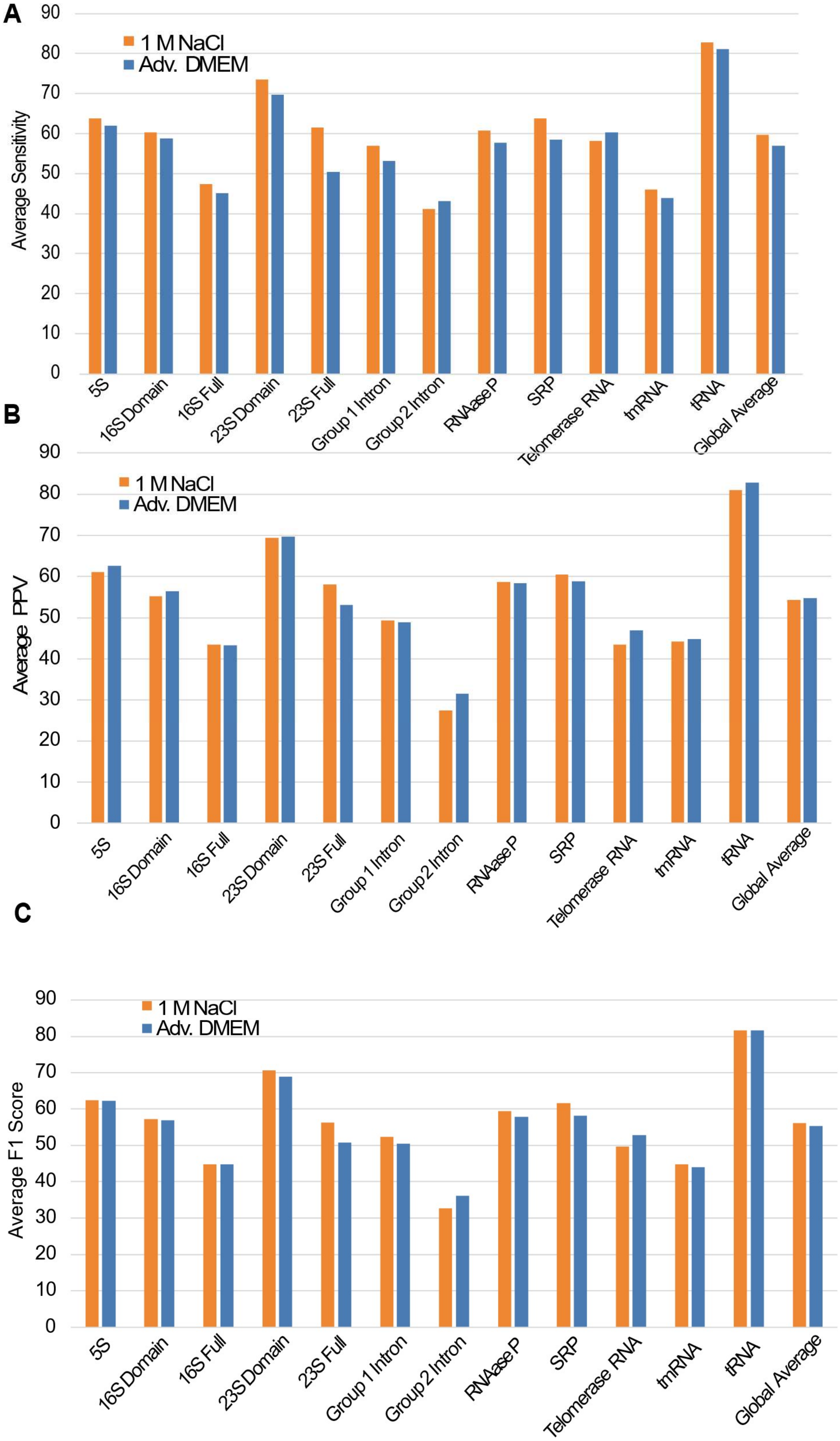

**Supplementary Figure 4. Comparison of MEA secondary structure prediction accuracy using standard 1 M NaCl and Adv. DMEM nearest neighbor parameters. (A)** Average sensitivity scores of maximum expected accuracy (MEA) RNA secondary structure predictions across ten ncRNA families using standard 1 M NaCl nearest neighbor parameters (orange) and Adv. DMEM-derived parameters (blue). For 16S and 23S rRNAs, predictions were evaluated separately for full-sequence structures and individual domain structures [3,4]. One-sided paired t-tests were performed to assess whether structure predictions improved using Adv. DMEM parameters; significant P-values are indicated.

**(B)** Average positive predictive value (PPV) scores of MEA RNA secondary structure predictions across the same ncRNA families and parameter sets described in **(A)**. One-sided paired t-tests were performed to assess whether structure predictions improved using Adv. DMEM parameters; significant P-values are indicated.

**(C)** Average F1 scores of MEA RNA secondary structure predictions across the same ncRNA families and parameter sets described in **(A)**. One-sided paired t-tests were performed to assess whether structure predictions improved using Adv. DMEM parameters; significant P-values are indicated.

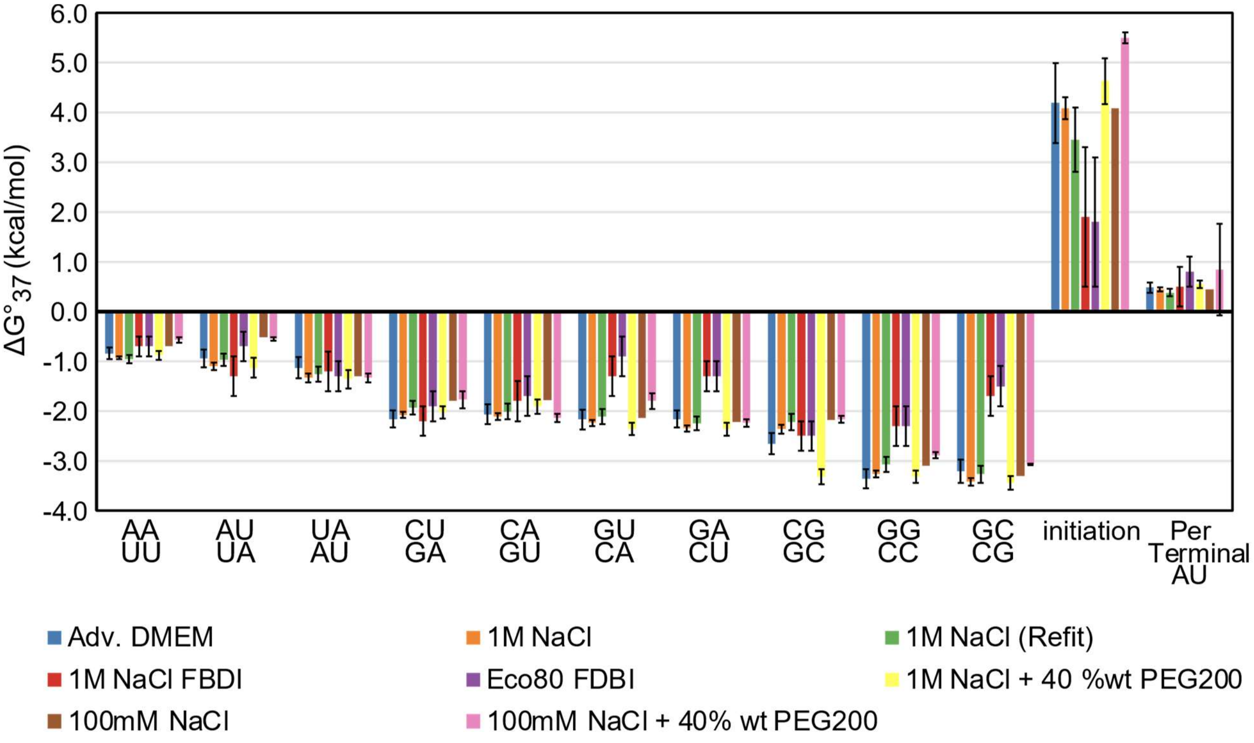

**Supplementary Figure 5. Comparison of WCF Nearest Neighbor Parameters in Adv. DMEM vs Other Models.** Nearest neighbor parameters for WCF base pair stacks, helix initiation, and A-U terminal ends were determined from folding stability of WCF helices measured via UV optical melting in Adv. DMEM of 42 helices (blue) and standard Turner 2004 parameters derived in 1 M NaCl of 90 helices (orange) [5]. WCF nearest neighbor parameters were recalculated for 40 helices measured in the Adv. DMEM dataset using the folding free energy measured in 1 M NaCl (green) to facilitate a direct comparison to the Adv. DMEM parameters [5]. The error bars represent the parameter's uncertainty estimate derived from the standard error of the regression values. Additional parameters determined from 24 helices using a fluorescence-detected binding isotherm (FDBI) assay are shown for measurements performed in 1 M NaCl (red) and Eco80 artificial cytoplasm (purple), an intracellular mimetic containing E. coli metabolites and ion concentrations from the Sieg et al. 2023 study [6]. Nearest neighbor parameters for helical stacks were previously derived in 1 M NaCl with 20 wt% PEG200 using 38 RNA helices in the Adams and Znosko 2019 study (yellow) [7]. The Adams and Znosko 2019 study included the symmetry term for self-complimentary helices as one of the independent parameters in their linear regression. The derived symmetry term (0.68 ± 0.21) overlaps with the expected symmetry correction term (0.43). Parameters determined from 45 helices using experimental UV melting measurements in 100 mM NaCl, 10 mM Na2HPO4, and 1 mM Na2EDTA with 40 wt% PEG200 (polyethylene glycol) as a synthetic crowder (pink) and without the crowding agent (brown) from the Ghosh et al. 2023 study. Error bars represent uncertainty estimates derived from the standard error of the regression values when available.

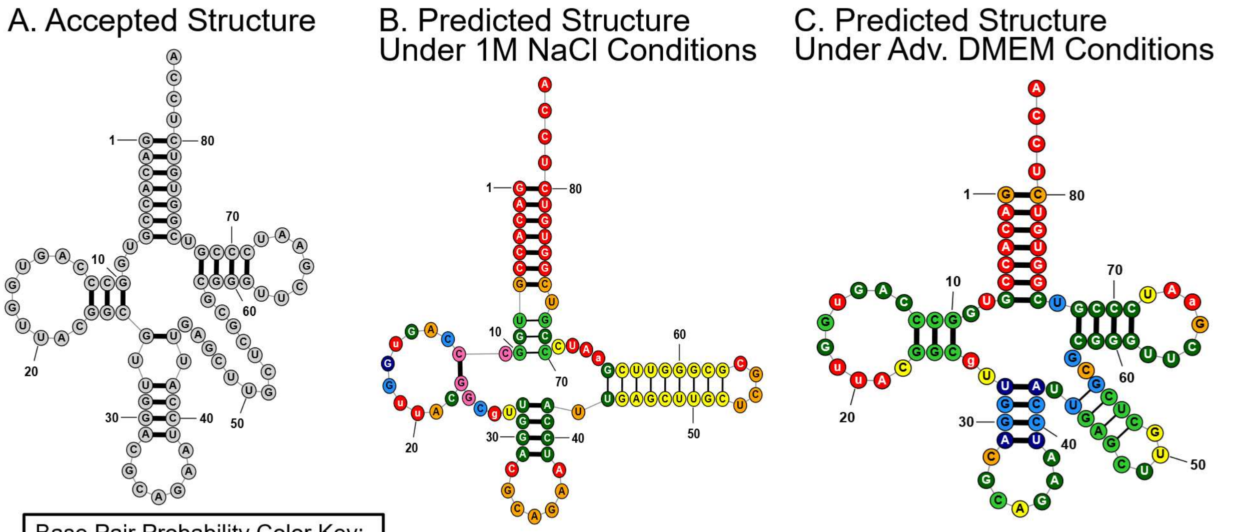

Base Pair Probability Color Key:

- P ≥ 0.99
- 0.99 > P ≥ 0.95
- 0.95 > P ≥ 0.90
- 0.90 > P ≥ 0.80
- 0.80 > P ≥ 0.70
- 0.70 > P ≥ 0.60
- 0.60 > P ≥ 0.50
- 0.50 > P

D. Comparison in Accuracy Metrics of Models

| Condition | Sensitivity | PPV | NED |
|---|---|---|---|
| 1M NaCl | 63.16 | 50.00 | 0.779 |
| Adv. DMEM | 94.74 | 81.82 | 0.391 |

**Supplementary Figure 6. Comparison of MEA structure predictions for a representative tRNA sequence, tRNA_tdbR00000409-Candida_cylindracea-44322-Ser-CAG, under Adv. DMEM and 1M NaCl nearest neighbor models. (A)** Accepted tRNA secondary structure based on comparative sequence analysis . **(B)** MEA predicted secondary structure using the 1 M NaCl nearest-neighbor model. **(C)** MEA predicted secondary structure using the Adv. DMEM model. **(D)** Table comparing sensitivity, positive predictive value (PPV), and normalized ensemble defect (NED) for the two model predictions for this tRNA sequence. All structures were generated with RNAstructure using the *MaxExpect* program. Base pairs matching the accepted comparative structure are shown with thick lines, whereas incorrectly predicted base pairs are shown with thin lines. Base pairs are additionally color-coded according to their computed pairing probabilities generated from the *partition* program in RNAstructure.

**Supplementary Table 1. Optical melting data for WCF RNA helices in Adv. DMEM.**
RNA helices are represented by a single sequence for self-complementary helices and by two sequences (5′–3′) for non-self-complementary helices. Values in the "Average of curve fits" columns represent the mean ± standard error of the thermodynamic parameters obtained from individual melting curve fits. Values in the "$T_M^{-1}$ vs. log $C_T$ plots" columns were obtained from linear regression of the reciprocal melting temperature ($T_M^{-1}$) as a function of the logarithm of total strand concentration ($C_T$). $\Delta G^{\circ}_{37}$ values were calculated from the corresponding $\Delta H^{\circ}$ and $\Delta S^{\circ}$ values at 37°C. Reported $T_M$ values are the melting temperatures calculated at 0.1 mM total oligonucleotide concentration.

| Duplexes 5' – 3' | Average of curve fits | | | | $T_M^{-1}$ vs log $C_T$ plots | | | |
|---|---|---|---|---|---|---|---|---|
| | -ΔH° (kcal/mol) | -ΔS° (eu) | -ΔG°37 (kcal/mol) | $T_M$ (°C) | -ΔH° (kcal/mol) | -ΔS° (eu) | -ΔG°37 (kcal/mol) | $T_M$ (°C) |
| AGAUAUCU | 59.5±5.7 | 175.2±18.3 | 5.17±0.10 | 34.4 | 54.8±3.6 | 159.8±11.6 | 5.19±0.06 | 34.3 |
| AUCUAGAU | 57.5±4.7 | 163.5±15.2 | 6.74±0.07 | 42.8 | 59.6±1.3 | 170.6±4.2 | 6.68±0.01 | 42.3 |
| AACUAGUU | 54.9±6.1 | 152.2±19.3 | 6.73±0.19 | 43.0 | 56.6±2.6 | 160.8±8.5 | 6.68±0.04 | 42.6 |
| AGUUAACU | 52.4±2.8 | 150.8±9.3 | 5.64±0.06 | 36.8 | 53.2±0.8 | 153.5±2.5 | 5.62±0.01 | 36.7 |
| ACUUAAGU | 54.7±6.2 | 157.8±20.1 | 5.74±0.16 | 37.3 | 52.9±2.0 | 152.1±6.7 | 5.69±0.04 | 37.1 |
| GAACGUUC | 66.9±1.8 | 189.5±5.6 | 8.14±0.09 | 48.8 | 71.6±3.2 | 204.1±10.0 | 8.25±0.09 | 48.5 |
| GUUCGAAC | 66.3±4.7 | 188.0±14.6 | 7.96±0.13 | 48.1 | 66.8±2.2 | 189.7±7.0 | 7.93±0.06 | 47.8 |
| UCAUAUGA | 54.3±2.1 | 158.0±6.5 | 5.32±0.10 | 35.0 | 60.1±2.7 | 178.7±8.9 | 5.25±0.04 | 34.8 |
| UAGAUCUA | 59.6 ±2.0 | 173.7± 6.4 | 5.66 ±0.09 | 36.9 | 63.7±1.9 | 187.3±6.1 | 5.60±0.03 | 36.6 |
| GUCGAC | 55.7±3.1 | 151.2±9.7 | 6.75±0.15 | 43.3 | 49.2±3.5 | 137.1±11.2 | 6.66±0.08 | 43.4 |
| GACGUC | 57.5±3.3 | 162.0±10.3 | 7.29±0.11 | 45.9 | 58.2±1.1 | 164.3±3.6 | 7.28±0.02 | 45.8 |
| ACUAUAGU | 59.4±4.4 | 171.0±14.2 | 6.35±0.04 | 40.6 | 60.5±1.7 | 174.8±5.5 | 6.30±0.02 | 40.2 |
| UGAUCA | 37.7±2.4 | 107.0±7.9 | 4.52±0.09 | 27.7 | 42.8±2.2 | 124.1±7.3 | 4.33±0.09 | 27.5 |
| GCAUGC | 58.5±7.1 | 166.8±22.8 | 6.79±0.16 | 43.0 | 51.8±2.4 | 145.5±7.8 | 6.70±0.04 | 43.2 |
| GUGCAC | 55.0±3.8 | 148.6±12.0 | 6.93±0.16 | 44.5 | 48.1±1.7 | 133.1±5.6 | 6.84±0.03 | 44.7 |
| AGCGCU | 50.3±5.3 | 137.1±16.3 | 7.74±0.23 | 50.3 | 44.6±2.0 | 119.5±6.4 | 7.49±0.07 | 50.2 |
| GUCUAGAC | 77.0±4.4 | 217.9±13.4 | 9.48±0.25 | 53.1 | 82.7±3.8 | 235.3±11.7 | 9.73±0.16 | 53.0 |
| GAUAUAUC | 63.6±6.9 | 187.8±22.2 | 5.53±0.05 | 35.3 | 58.3±2.1 | 170.5±6.8 | 5.38±0.05 | 35.5 |
| GUAUAUAC | 60.1±2.1 | 177.7±7.0 | 5.00±0.06 | 33.6 | 59.7±1.4 | 176.5±4.8 | 4.99±0.04 | 33.5 |
| UGCGCA | 50.9±5.3 | 138.0±16.4 | 8.09±0.19 | 52.5 | 49.4±1.4 | 133.5±4.4 | 7.94±0.05 | 52.1 |
| AUACGUAU | 56.2±1.6 | 162.2±4.9 | 5.87±0.10 | 38.1 | 57.0±3.6 | 165.0±11.8 | 5.86±0.07 | 38.0 |
| CACGUG | 53.1±4.2 | 149.7±13.4 | 6.60±0.11 | 42.5 | 50.4±1.3 | 141.4±4.1 | 6.53±0.01 | 42.4 |

| | | | | | | | | | |
|---|---|---|---|---|---|---|---|---|---|
| CCAUGG | 55.9±3.2 | 157.2±10.3 | 7.12±0.11 | 45.2 | | 61.2±2.1 | 174.1±6.6 | 7.16±0.03 | 44.7 |
| CCGCGG | 65.3±1.9 | 176.6±5.7 | 10.48±0.14 | 61.6 | | 69.4±1.9 | 189.1±5.7 | 10.75±0.13 | 61.5 |
| CCUAGG | 57.3±2.2 | 159.3±7.0 | 7.87±0.10 | 49.4 | | 60.7±1.8 | 170.2±5.5 | 7.94 ±0.05 | 49.0 |
| GAGCUC | 57.3±4.9 | 161.0±15.4 | 7.35±0.16 | 46.3 | | 58.7±4.6 | 165.7±14.6 | 7.35±0.13 | 46.1 |
| GCUAGC | 56.0±5.6 | 155.5±17.4 | 7.74±0.18 | 48.8 | | 52.8±2.2 | 145.8±6.8 | 7.59±0.06 | 48.6 |
| UCGCGA | 49.5±2.1 | 137.0±6.5 | 6.97±0.07 | 45.3 | | 51.2±0.8 | 142.6±2.4 | 6.97±0.01 | 45.1 |
| UCUAUAGA | 61.8±1.8 | 179.6±5.9 | 6.13±0.12 | 39.3 | | 65.9±3.0 | 192.9±9.8 | 6.09±0.04 | 38.9 |
| UUCCGGAA | 67.5±3.2 | 183.9±9.6 | 10.46±0.26 | 60.7 | | 71.4±3.3 | 195.6±10.0 | 10.71±0.20 | 60.5 |
| UUGCGCAA | 56.0±2.1 | 150.9±6.6 | 9.13±0.15 | 57.4 | | 63.5±3.3 | 174.2±10.0 | 9.49±0.16 | 56.8 |
| UUGUACAA | 50.8±0.9 | 144.5±3.0 | 6.01±0.09 | 39.1 | | 56.1±1.6 | 161.7±5.1 | 5.97±0.02 | 38.6 |
| UUGGCCAA | 59.8±4.2 | 160.4±12.7 | 10.07±0.26 | 61.6 | | 60.8±2.9 | 163.4±8.6 | 10.11±0.18 | 61.4 |
| 5'GGCUUCAA<br>3'CCGAAGUU | 66.7±2.6 | 184.0±7.8 | 9.81±0.17 | 53.0 | | 67.0±2.6 | 184.5±7.9 | 9.80±0.14 | 52.9 |
| 5'AAGGUUGGAA<br>3'UUCCAACCUU | 81.4±3.8 | 222.3±11.5 | 12.41±0.23 | 61.2 | | 76.9±1.2 | 209.0±3.6 | 12.11±0.10 | 61.3 |
| GGAUCC | 54.8±3.6 | 154.0±11.2 | 7.00±0.16 | 44.7 | | 49.0±1.5 | 135.7±4.7 | 6.86±0.03 | 44.7 |
| GGUACC | 55.4±4.3 | 155.8±13.5 | 7.04±0.16 | 44.8 | | 57.1±2.5 | 161.6±8.0 | 7.02±0.04 | 44.5 |
| CGCGCG | 57.9±5.4 | 156.6±16.3 | 9.32±0.35 | 57.8 | | 54.1±3.2 | 145.1±9.6 | 9.07±0.18 | 57.7 |
| GCGCGC | 64.6±3.7 | 174.3±10.9 | 10.53±0.28 | 62.2 | | 61.3±1.3 | 164.5±3.8 | 10.29± 0.09 | 62.2 |

**Supplementary Table 2. Comparison of WCF nearest neighbor parameters.**
Nearest neighbor free energy parameters ($\Delta G^{\circ}_{37}$) for WCF RNA helices derived in Adv. DMEM and 1 M NaCl are shown for comparison [5]. Parameters include the eight WCF nearest-neighbor stacks, the initiation term, and the terminal A–U pair contribution. σ represents the uncertainty associated with each derived parameter.

| WCF Stack: | Adv. DMEM $\Delta G^{\circ}_{37}$ (kcal/mol): | σ (kcal/mol): | 1M NaCl $\Delta G^{\circ}_{37}$ (kcal/mol): | σ (kcal/mol): |
|---|---|---|---|---|
| AA<br>UU | -0.84 | 0.12 | -0.93 | 0.03 |
| AU<br>UA | -0.94 | 0.18 | -1.10 | 0.08 |
| UA<br>AU | -1.13 | 0.21 | -1.33 | 0.09 |
| CU<br>GA | -2.16 | 0.17 | -2.08 | 0.06 |
| CA<br>GU | -2.06 | 0.20 | -2.11 | 0.07 |
| GU<br>CA | -2.17 | 0.20 | -2.24 | 0.06 |
| GA<br>CU | -2.16 | 0.17 | -2.35 | 0.06 |
| CG<br>GC | -2.65 | 0.22 | -2.36 | 0.09 |
| GG<br>CC | -3.36 | 0.20 | -3.26 | 0.07 |
| GC<br>CG | -3.21 | 0.23 | -3.42 | 0.08 |
| Initiation | 4.19 | 0.81 | 4.09 | 0.22 |
| Terminal A-U Pair | 0.48 | 0.11 | 0.45 | 0.04 |

**Supplementary Table 3. Optical melting data for G-U-containing RNA helices in Adv. DMEM.**

RNA helices are represented by a single sequence for self-complementary helices and by two sequences (5′–3′) for non-self-complementary helices. Thermodynamic parameters were determined by UV optical melting under Adv. DMEM conditions. Values in the "Average of curve fits" columns represent the mean ± standard error of the thermodynamic parameters obtained from individual melting curve fits. Values in the "$T_M^{-1}$ vs. log $C_T$ plots" columns were obtained from linear regression of the reciprocal melting temperature ($T_M^{-1}$) as a function of the logarithm of total strand concentration ($C_T$). $\Delta G^\circ_{37}$ values were calculated from the corresponding $\Delta H^\circ$ and $\Delta S^\circ$ values at 37˚C. Reported $T_M$ values are the melting temperatures calculated at 0.1 mM total oligonucleotide concentration.

| Duplexes (5'-3') | Average of curve fits | | | | | $T_M^{-1}$ vs log $C_T$ plots | | | |
|---|---|---|---|---|---|---|---|---|---|
| | -ΔH° (kcal/mol) | -ΔS° (eu) | $-\Delta G^\circ_{37}$ (kcal/mol) | $T_M$[b] (˚C) | | -ΔH° (kcal/mol) | -ΔS° (eu) | $-\Delta G^\circ_{37}$ (kcal/mol) | $T_M$[b] (˚C) |
| GGCGCU | 53.5±1.9 | 146.8±5.9 | 7.95±0.10 | 50.8 | | 54.1±2.6 | 148.7±8.0 | 7.96±0.10 | 50.6 |
| GACGCGUU | 65.0±3.5 | 179.9±10.9 | 9.21±0.19 | 54.8 | | 67.2±4.7 | 186.8±14.5 | 9.32±0.23 | 54.8 |
| GAGUGCUC | 77.8±2.1 | 224.3±6.5 | 8.28±0.11 | 47.7 | | 77.2±2.4 | 222.3±7.7 | 8.25±0.07 | 47.7 |
| GCAGCUGU | 69.2±2.5 | 192.5±7.6 | 9.50±0.20 | 55.1 | | 74.5±5.0 | 208.8±15.3 | 9.76±0.26 | 55.0 |
| GGCGUGCC | 70.8±4.2 | 199.0±13.2 | 9.08±0.10 | 52.6 | | 73.0±4.4 | 205.9±13.7 | 9.16±0.18 | 52.5 |
| GGCUGGCC | 86.6±2.3 | 239.0±6.8 | 12.46±0.22 | 63.4 | | 87.6±5.1 | 242.1±15.2 | 12.53±0.38 | 63.3 |
| UACCGGUG | 62.7±2.4 | 169.9±7.1 | 9.97±0.21 | 59.8 | | 59.3±5.2 | 159.7±15.6 | 9.76±0.35 | 59.9 |
| UCACGUGG | 64.5±2.5 | 180.0±7.8 | 8.64±0.10 | 52.0 | | 67.0±1.3 | 187.9±4.1 | 8.72±0.05 | 51.8 |
| UGACGUCG | 67.6±2.1 | 186.2±6.5 | 9.88±0.11 | 57.6 | | 73.6±6.7 | 204.5±20.6 | 10.19±0.38 | 57.3 |
| CCAGCGUCCU<br>GGUUGUAGGA | 101.4±1.7 | 288.7±5.5 | 11.90±0.10 | 54.3 | | 99.7±4.3 | 283.3±13.2 | 11.81±0.21 | 54.3 |
| CGGUGCAUCG | 93.7±3.2 | 262.4±9.6 | 12.32±0.2 | 60.7 | | 93.6±5.2 | 262.2±15.8 | 12.31±0.33 | 60.6 |
| CAGAGGAGAC<br>GUCUUUUCUG | 94.0±5.7 | 275.0±17.9 | 8.68±0.15 | 44.3 | | 89.3±3.1 | 260.3±10.0 | 8.57±0.06 | 44.2 |
| GAGGUGAG<br>CUCUGCUC | 71.6±1.7 | 210.21±5.4 | 6.37±0.08 | 36.3 | | 64.5±1.2 | 187.2±4.0 | 6.42±0.01 | 36.5 |
| GUGUGCAUAC | 77.2±4.7 | 221.5±14.9 | 8.45±0.12 | 48.6 | | 82.7±3.8 | 238.8±10.6 | 8.59±0.09 | 48.3 |
| GAGUGGAGAG<br>CUCAUUUCUC | 88.4±5.3 | 257.0±16.9 | 8.73±0.09 | 44.9 | | 87.1±2.0 | 252.9±6.4 | 8.68±0.04 | 44.8 |

| | | | | | | | | | |
|---|---|---|---|---|---|---|---|---|---|
| CAGCGCGUUG | 60.5±2.8 | 164.1±8.1 | 9.60±0.26 | 58.5 | | 71.8±4.4 | 198.4±13.3 | 10.26±0.26 | 58.2 |
| GAUGCAUU | 57.5±1.1 | 167.2±3.9 | 5.69±0.11 | 37.1 | | 58.8±3.5 | 171.2±11.4 | 5.69±0.06 | 37.1 |
| GGACGUGUCC | 87.1±4.4 | 247.5±13.4 | 10.28±0.28 | 54.3 | | 81.9±4.4 | 231.8±13.5 | 10.01±0.21 | 54.3 |
| CUCGGCUC GAGUUGAG | 76.6±2.6 | 223.4±8.3 | 7.30±0.05 | 40.1 | | 75.9±1.9 | 221.3±6.1 | 7.28±0.03 | 40.1 |
| GAGUGGAG CUCGUCUC | 83.1±2.2 | 240.0±6.9 | 8.68±0.09 | 45.2 | | 80.6±1.8 | 232.1±5.7 | 8.59±0.06 | 45.1 |
| GAGAGCUUUC | 76.0±4.9 | 219.5±15.8 | 7.92±0.09 | 46.4 | | 81.0±3.5 | 235.3±11.1 | 7.98±0.07 | 46.1 |
| GCUGGUGC CGAUUACG | 67.0±1.6 | 193.6±5.1 | 6.92±0.08 | 38.8 | | 63.0±2.6 | 180.9±8.3 | 6.87±0.04 | 38.7 |
| GAGUGGAGAG CUCAUUUCUC | 92.4±2.9 | 269.5±9.2 | 8.82±0.10 | 44.9 | | 90.4±3.8 | 263.1±12.0 | 8.78±0.10 | 44.9 |
| CCUGUAGG | 58.2±1.3 | 170.4±4.4. | 5.37±0.09 | 35.4 | | 62.7±3.2 | 185.2±10.7 | 5.27±0.08 | 35.0 |
| GGAGUUCC | 68.3±3.0 | 201.9±9.6 | 5.65±0.09 | 36.9 | | 68.9±4.0 | 204.2±13.0 | 5.63±0.07 | 36.8 |

**Supplementary Table 4. Comparison of G-U stack nearest neighbor parameters.**
Nearest-neighbor free-energy parameters ($\Delta G^{\circ}_{37}$) for G-U containing RNA helices derived in Adv. DMEM and 1 M NaCl are shown for comparison [8]. Parameters include the eleven G-U nearest-neighbor stacks. σ represents the uncertainty associated with each derived parameter.

| G-U Stack: | Adv. DMEM $\Delta G^{\circ}_{37}$ (kcal/mol): | σ (kcal/mol): | 1M NaCl $\Delta G^{\circ}_{37}$ (kcal/mol): | σ (kcal/mol): |
|---|---|---|---|---|
| GC<br>UG | -1.35 | 0.55 | -2.15 | 0.10 |
| GG<br>UC | -1.58 | 0.67 | -1.77 | 0.09 |
| GG<br>CU | -1.21 | 0.61 | -1.80 | 0.09 |
| GA<br>UU | -0.40 | 0.48 | -0.51 | 0.08 |
| GU<br>UA | -0.72 | 0.67 | -0.9 | 0.08 |
| CG<br>GU | -1.27 | 0.56 | -1.25 | 0.09 |
| UG<br>AU | -0.24 | 0.55 | -0.39 | 0.09 |
| AG<br>UU | -0.11 | 0.55 | -0.35 | 0.08 |
| UG<br>GU | 0.32 | 1.26 | -0.57 | 0.19 |
| GU<br>UG | 2.08 | 1.03 | 0.72 | 0.19 |
| GG<br>UU | 0.47 | 0.83 | -0.25 | 0.16 |

**Supplementary Table 5. Optical melting data for RNA duplexes containing dangling ends.** RNA helices are represented by a single sequence for self-complementary duplexes. The entries marked with * are the reference duplexes without any dangling ends. Thermodynamic parameters were determined by UV optical melting under Adv. DMEM conditions. Values in the "Average of curve fits" columns represent the mean ± standard error of the thermodynamic parameters obtained from individual melting curve fits. Values in the "$T_M^{-1}$ vs. log $C_T$ plots" columns were obtained from linear regression of the reciprocal melting temperature ($T_M^{-1}$) as a function of the logarithm of total strand concentration ($C_T$). $\Delta G^\circ_{37}$ values were calculated from the corresponding $\Delta H^\circ$ and $\Delta S^\circ$ values at 37°C. Reported $T_M$ values are the melting temperatures calculated at 0.1 mM total oligonucleotide concentration.

| Duplexes 5' – 3' | Average of curve fits | | | | $T_M^{-1}$ vs log $C_T$ plots | | | |
|---|---|---|---|---|---|---|---|---|
| | -ΔH° (kcal/mol) | -ΔS° (eu) | $-\Delta G^\circ_{37}$ (kcal/mol) | $T_M$ (°C) | -ΔH° (kcal/mol) | -ΔS° (eu) | $-\Delta G^\circ_{37}$ (kcal/mol) | $T_M$ (°C) |
| UUGCGCAA* | 59.2±2.5 | 160.6±7.8 | 9.34±0.17 | 57.5 | 65.0±4.9 | 178.5±14.9 | 9.66±0.3 | 57.2 |
| AUUGCGCAA | 66.3±12.8 | 181.0±39.9 | 10.15±0.71 | 59.5 | 75.5±0.9 | 208.9±2.7 | 10.74±0.05 | 59.3 |
| UUUGCGCAA | 59.3±1.6 | 160.4±4.6 | 9.52±0.21 | 58.5 | 68.9±4.7 | 177.5±14.5 | 9.80±0.24 | 58.1 |
| CUUGCGCAA | 62.4±1.9 | 167.0±5.8 | 9.68±0.13 | 58.3 | 68.3±4.7 | 191.1±8.3 | 10.02±0.14 | 57.8 |
| GUUGCGCAA | 63.2±6.4 | 172.1±19.6 | 9.83±0.34 | 58.8 | 68.5±2.3 | 188.3±7.0 | 10.15±0.12 | 58.6 |
| AUGCGCAU* | 59.8±1.9 | 161.2±5.6 | 9.76±0.17 | 59.8 | 60.4±3.4 | 163.2±10.3 | 9.81±0.2 | 59.8 |
| AAUGCGCAU | 61.6±2.6 | 166.4±7.6 | 9.96±0.24 | 60.2 | 69.7±2.9 | 191.1±8.7 | 10.43±0.16 | 59.7 |
| UAUGCGCAU | 61.5±2.4 | 166.6±7.0 | 9.84±0.20 | 59.5 | 67.1±2.3 | 183.7±7.2 | 10.17±0.13 | 59.2 |
| CAUGCGCAU | 60.5±3.1 | 165.1±9.6 | 9.27±0.13 | 56.6 | 58.8±2.9 | 159.8±8.9 | 9.20±0.17 | 56.8 |
| GAUGCGCAU | 60.2±4.3 | 164.0±13.1 | 9.30±0.33 | 56.9 | 63.6±4.5 | 174.2±13.5 | 9.54±0.28 | 57.1 |
| CUGCAG* | 55.3±5.8 | 156.3±18.7 | 6.84±0.12 | 43.7 | 53.2±1.9 | 149.4±6.1 | 6.84±0.03 | 43.9 |
| ACUGCAG | 58.0±2.4 | 162.0±7.6 | 7.83±0.10 | 49.0 | 64.6±1.0 | 182.3±3.0 | 8.03±0.03 | 48.7 |
| UCUGCAG | 57.2±1.4 | 161.1±4.0 | 7.25±0.10 | 45.7 | 59.5±2.5 | 168.2±7.9 | 7.30±0.06 | 45.7 |
| CCUGCAG | 59.0±1.8 | 164.5±3.8 | 7.93±0.05 | 49.4 | 63.7±1.0 | 179.3±3.1 | 8.08±0.03 | 49.1 |
| GCUGCAG | 55.5±1.8 | 155.1±5.4 | 7.42±0.13 | 47.1 | 59.0±3.8 | 165.9±11.8 | 7.52±0.12 | 47.0 |

| | | | | | | | | | |
|---|---|---|---|---|---|---|---|---|---|
| GUGCAC* | 55.0±3.8 | 148.6±12.0 | 6.93±0.16 | 44.5 | | 48.1±1.7 | 133.1±5.6 | 6.84±0.03 | 44.7 |
| AGUGCAC | 56.7±3.1 | 157.8±9.4 | 7.68±0.20 | 48.3 | | 63.5±3.8 | 179.3±11.8 | 7.87±0.12 | 48.1 |
| UGUGCAC | 54.9±1.4 | 153.8±4.3 | 7.19±0.11 | 45.8 | | 58.4±2.6 | 165.0±8.1 | 7.26±0.06 | 45.6 |
| CGUGCAC | 57.9±1.6 | 162.0±5.1 | 7.66±0.08 | 48.0 | | 61.8±2.0 | 174.0±6.1 | 7.77±0.06 | 47.9 |
| GGUGCAC | 58.5±2.6 | 163.5±8.1 | 7.77±0.13 | 48.5 | | 60.2±2.7 | 168.9±8.6 | 7.81±0.08 | 48.4 |
| UUGCGCAAA | 63.2±5.3 | 171.2±15.6 | 10.10±0.41 | 60.4 | | 70.5±2.1 | 193.3±6.4 | 10.57±0.12 | 60.1 |
| UUGCGCAAU | 64.9±2.8 | 176.9±8.3 | 10.05±0.21 | 59.4 | | 69.2±1.1 | 189.8±3.3 | 10.31±0.06 | 59.2 |
| UUGCGCAAC | 64.5±2.2 | 175.0±6.4 | 10.19±0.21 | 60.4 | | 68.5±4.1 | 187.1±12.3 | 10.44±0.25 | 60.2 |
| UUGCGCAAG | 65.3±3.1 | 171.8±9.4 | 10.02±0.19 | 59.9 | | 65.9±4.1 | 179.8±12.6 | 10.20±0.24 | 59.7 |
| AUGCGCAUA | 64.3±3.9 | 174.0±11.4 | 10.33±0.37 | 61.2 | | 79.3±3.0 | 219.3±9.1 | 11.23±0.12 | 60.4 |
| AUGCGCAUU | 61.5±3.6 | 166.2±10.6 | 9.96±0.29 | 60.2 | | 64.7±3.3 | 176.0±10.1 | 10.16±0.19 | 60.1 |
| AUGCGCAUC | 62.8±2.5 | 170.1±7.3 | 10.06±0.20 | 60.3 | | 64.8±2.7 | 176.1±8.3 | 10.21±0.17 | 60.3 |
| AUGCGCAUG | 65.8±9.1 | 179.4±27.3 | 10.12±0.63 | 59.5 | | 76.8±1.9 | 212.7±5.7 | 10.82±0.11 | 59.3 |
| CUGCAGA | 65.1±1.1 | 181.1±3.4 | 8.91±0.11 | 53.2 | | 71.5±2.6 | 200.8±7.9 | 9.17±0.10 | 52.9 |
| CUGCAGU | 56.6±1.9 | 157.7±6.0 | 7.71±0.13 | 48.6 | | 63.0±3.3 | 177.8±10.2 | 7.86±0.09 | 48.1 |
| CUGCAGC | 54.0±3.0 | 150.1±9.3 | 7.45±0.19 | 47.5 | | 50.1±5.9 | 137.9±18.4 | 7.32±0.24 | 47.5 |
| CUGCAGG | 67.4±4.2 | 188.7±13.0 | 8.86±0.17 | 52.4 | | 70.8±1.7 | 199.7±5.2 | 9.03±0.06 | 52.4 |
| GUGCACA | 72.1±4.3 | 198.4±12.8 | 10.55±0.29 | 59.5 | | 79.5±2.5 | 220.7±7.5 | 11.00±0.14 | 59.3 |
| GUGCACU | 66.6±4.3 | 184.9±13.0 | 9.22±0.22 | 54.5 | | 68.2±2.1 | 189.7±6.6 | 9.23±0.01 | 54.5 |
| GUGCACC | 59.4±2.4 | 163.4±7.2 | 8.69±0.17 | 53.6 | | 64.4±5.3 | 179.0±16.4 | 8.90±0.25 | 53.4 |
| GUGCACG | 62.0±2.4 | 171.8±7.3 | 8.75±0.17 | 53.2 | | 69.2±1.2 | 193.9±3.7 | 9.03±0.05 | 52.8 |

**Supplementary Table 6. Comparison of nearest neighbor parameters for dangling ends.** Nearest neighbor free energy parameters ($\Delta G^{\circ}_{37}$) for dangling ends derived in Adv. DMEM and 1 M NaCl are shown for comparison [2]. Parameters include 16 3' dangling end and 16 5' dangling end combinations. σ represents the uncertainty associated with each derived parameter.

| Parameter 5' --> 3' | Adv. DMEM $\Delta G^{\circ}_{37}$ (kcal/mol): | σ (kcal/mol): | 1M NaCl $\Delta G^{\circ}_{37}$ (kcal/mol): | σ (kcal/mol): |
|---|---|---|---|---|
| AU<br>A | -0.54 | 0.58 | -0.30 | 0.29 |
| UU<br>A | -0.07 | 0.55 | -0.22 | 0.33 |
| CU<br>A | -0.18 | 0.56 | -0.14 | 0.15 |
| GU<br>A | -0.25 | 0.56 | -0.22 | 0.33 |
| AA<br>U | -0.31 | 0.57 | -0.33 | 0.12 |
| UA<br>U | -0.18 | 0.57 | -0.19 | 0.12 |
| CA<br>U | 0.31 | 0.54 | -0.25 | 0.12 |
| GA<br>U | 0.14 | 0.55 | -0.35 | 0.12 |
| AC<br>G | -0.60 | 0.42 | -0.53 | 0.13 |
| UC<br>G | -0.23 | 0.40 | -0.15 | 0.11 |
| CC<br>G | -0.62 | 0.42 | -0.36 | 0.17 |
| GC<br>G | -0.34 | 0.41 | -0.17 | 0.13 |
| AG<br>C | -0.52 | 0.42 | -0.22 | 0.16 |
| UG<br>C | -0.21 | 0.40 | -0.07 | 0.10 |
| CG<br>C | -0.470 | 0.41 | -0.27 | 0.12 |

| | | | | |
|---|---|---|---|---|
| GG<br>C | -0.49 | 0.42 | 0.04 | 0.12 |
| AA<br>U | -0.46 | 0.57 | -0.74 | 0.29 |
| AU<br>U | -0.33 | 0.57 | -0.58 | 0.30 |
| AC<br>U | -0.39 | 0.57 | -0.49 | 0.30 |
| AG<br>U | -0.27 | 0.56 | -0.83 | 0.29 |
| UA<br>A | -0.71 | 0.60 | -0.66 | 0.12 |
| UU<br>A | -0.18 | 0.57 | -0.09 | 0.11 |
| UC<br>A | -0.20 | 0.57 | -0.13 | 0.10 |
| UG<br>A | -0.50 | 0.58 | -0.66 | 0.12 |
| GA<br>C | -1.17 | 0.46 | -1.15 | 0.11 |
| GU<br>C | -0.51 | 0.42 | -0.62 | 0.09 |
| GC<br>C | -0.24 | 0.40 | -0.35 | 0.11 |
| GG<br>C | -1.10 | 0.45 | -1.25 | 0.16 |
| CA<br>G | -2.08 | 0.52 | -1.73 | 0.17 |
| CU<br>G | -1.20 | 0.46 | -1.19 | 0.13 |
| CC<br>G | -1.03 | 0.45 | -0.78 | 0.13 |
| CG<br>G | -1.10 | 0.45 | -1.64 | 0.16 |

**Supplementary Table 7. Thermodynamic parameters of RNA duplexes containing terminal mismatches.**

RNA helices are represented by a single sequence for self-complementary duplexes. Thermodynamic parameters were determined by UV optical melting under Adv. DMEM conditions. Values in the "Average of curve fits" columns represent the mean ± standard error of the thermodynamic parameters obtained from individual melting curve fits. Values in the "$T_M^{-1}$ vs. log $C_T$ plots" columns were obtained from linear regression of the reciprocal melting temperature ($T_M^{-1}$) as a function of the logarithm of total strand concentration ($C_T$). $\Delta G^{\circ}_{37}$ values were calculated from the corresponding $\Delta H^{\circ}$ and $\Delta S^{\circ}$ values at 37°C. Reported $T_M$ values are the melting temperatures calculated at 0.1 mM total oligonucleotide concentration.

| Duplexes (5'-3') | Average of curve fits | | | | $T_M^{-1}$ vs log $C_T$ plots | | | |
|---|---|---|---|---|---|---|---|---|
| | -ΔH° (kcal/mol) | -ΔS° (eu) | $-\Delta G^{\circ}_{37}$ (kcal/mol) | $T_M$ (°C) | -ΔH° (kcal/mol) | -ΔS° (eu) | $-\Delta G^{\circ}_{37}$ (kcal/mol) | $T_M$ (°C) |
| AUUGCGCAAA | 63.1±1.7 | 170.1±5.4 | 10.38±0.07 | 62.0 | 71.7±3.6 | 196.1±10.9 | 10.89±0.23 | 61.3 |
| CUUGCGCAAC | 63.8±2.5 | 172.8±7.5 | 10.22±0.17 | 60.8 | 67.9±3.9 | 185.1±11.7 | 10.49±0.25 | 60.6 |
| GUUGCGCAAG | 65.2±2.2 | 177.0±7.0 | 10.33±0.10 | 60.8 | 68.6±6.2 | 187.2±18.7 | 10.53±0.39 | 60.6 |
| UUUGCGCAAU | 64.9±3.4 | 176.4±10.1 | 10.14±0.29 | 59.9 | 62.7±3.8 | 169.11.7 | 10.01±0.23 | 60.0 |
| ACUGCAGA | 61.9±6.0 | 170.2±18.6 | 9.07±0.22 | 55.0 | 63.9±3.2 | 176.5±9.8 | 9.13±0.14 | 54.7 |
| CCUGCAGC | 63.1±1.8 | 174.6±5.6 | 8.91±0.11 | 53.8 | 63.8±2.6 | 176.9±7.9 | 8.95±0.12 | 53.8 |
| GCUGCAGG | 68.3±2.3 | 189.3±7.3 | 9.55±0.10 | 55.7 | 70.2±3.0 | 195.3±9.1 | 9.64±0.16 | 55.6 |
| UCUGCAGU | 60.4±2.5 | 167.7±8.0 | 8.38±0.15 | 51.6 | 69.5±4.1 | 196.2±12.7 | 8.69±0.15 | 51.1 |
| CCUGCAGA | 63.9±1.8 | 176.5±5.6 | 9.12±0.11 | 54.7 | 63.1±1.8 | 174.3±5.5 | 9.07±0.08 | 54.6 |
| ACUGCAGC | 61.8±5.0 | 170.0±15.2 | 9.04±0.25 | 54.8 | 63.9±2.6 | 176.8±8.1 | 9.11±0.12 | 54.6 |
| ACUGCAGG | 64.6±2.7 | 177.9±8.3 | 9.42±0.16 | 56.1 | 66.4±1.7 | 183.6±5.2 | 9.50±0.08 | 55.9 |
| GCUGCAGA | 67.5±1.9 | 187.3±5.9 | 9.41±0.13 | 55.2 | 69.6±3.4 | 193.7210.5 | 9.52±0.18 | 55.1 |
| CCUGCAGU | 63.7±2.2 | 176.2±6.9 | 9.03±0.10 | 54.3 | 63.0±3.1 | 174.1±9.7 | 8.99±0.14 | 54.2 |
| AAUGCGCAUA | 64.7±2.7 | 174.6±8.1 | 10.55±0.23 | 62.3 | 68.9±3.7 | 187.0±11.2 | 10.86±0.28 | 62.3 |
| CAUGCGCAUC | 64.7±2.7 | 174.7±8.0 | 10.55±0.18 | 62.3 | 63.9±3.9 | 172.3±11.9 | 10.49±0.25 | 62.3 |
| AGUGCACA | 69.8±5.1 | 191.0±15.3 | 10.55±0.38 | 60.3 | 69.4±3.1 | 189.9±9.3 | 10.49±0.19 | 60.1 |
| CGUGCACC | 65.0±1.6 | 178.8±5.0 | 9.58±0.05 | 56.8 | 72.8±2.5 | 202.5±7.5 | 9.98±0.13 | 56.5 |

| | | | | | | | | | |
|---|---|---|---|---|---|---|---|---|---|
| AGACGCGUUA | 69.3±2.0 | 190.5±5.9 | 10.19±0.15 | 58.6 | | 72.9±2.1 | 201.6±6.5 | 10.39±0.11 | 58.4 |
| CGACGCGUUC | 73.3±1.3 | 203.2±4.0 | 10.31±0.13 | 57.9 | | 77.3±5.4 | 215.3±16.4 | 10.53±0.30 | 57.8 |
| GGACGCGUUG | 64.7±5.2 | 178.1±15.7 | 9.42±0.36 | 56.0 | | 74.5±3.7 | 208.1±11.3 | 9.91±0.18 | 55.7 |
| UGACGCGUUU | 67.8±2.5 | 187.8±7.7 | 9.21±0.20 | 55.6 | | 63.3±6.0 | 174.2±18.4 | 9.30±0.33 | 55.8 |

**Supplementary Table 8. Comparison of nearest neighbor parameters for terminal mismatches.** Nearest neighbor free energy parameters ($\Delta G^{\circ}_{37}$) for G-U containing RNA helices derived in Adv. DMEM and 1 M NaCl are shown for comparison [2]. Parameters include 21 unique terminal mismatch stacks. σ represents the uncertainty associated with each derived parameter.

| Parameter 5' --> 3' | Adv. DMEM $\Delta G^{\circ}_{37}$ (kcal/mol): | σ (kcal/mol): | 1M NaCl $\Delta G^{\circ}_{37}$ (kcal/mol): | σ (kcal/mol): |
|---|---|---|---|---|
| AA<br>UA | -0.62 | 0.58 | -0.75 | 0.30 |
| AC<br>UC | -0.42 | 0.57 | -0.63 | 0.34 |
| AG<br>UG | -0.44 | 0.57 | -0.85 | 0.22 |
| AU<br>UU | -0.18 | 0.56 | -0.79 | 0.45 |
| GA<br>CA | -1.15 | 0.46 | -1.09 | 0.16 |
| GC<br>CC | -1.06 | 0.45 | -0.71 | 0.16 |
| GG<br>CG | -1.40 | 0.47 | -1.47 | 0.19 |
| GU<br>CU | -0.93 | 0.44 | -0.77 | 0.15 |
| GA<br>CC | -1.12 | 0.45 | -0.88 | 0.20 |
| GC<br>CA | -1.14 | 0.46 | -1.09 | 0.14 |
| GG<br>CA | -1.33 | 0.47 | -1.58 | 0.19 |
| GA<br>CG | -1.34 | 0.47 | -1.24 | 0.17 |
| GU<br>CC | -1.08 | 0.45 | -0.98 | 0.19 |
| UA<br>AA | -0.53 | 0.59 | -0.94 | 0.13 |
| UC<br>AC | -0.34 | 0.57 | -0.59 | 0.11 |

| | | | | |
|---|---|---|---|---|
| CA<br>GA | -1.83 | 0.50 | -1.5 | 0.20 |
| CC<br>GC | -1.57 | 0.48 | -1.05 | 0.18 |
| UA<br>GA | -0.54 | 0.56 | -1.27 | 0.26 |
| UC<br>GC | -0.61 | 0.56 | -0.38 | 0.16 |
| UG<br>GG | -0.30 | 0.54 | -0.81 | 0.35 |
| UU<br>GU | 0.01 | 0.53 | -0.4 | 0.32 |

**Supplementary Table 9. Optical melting experiment results for RNA duplexes containing hairpins.**

Thermodynamic parameters for RNA hairpins were determined by UV optical melting under Adv. DMEM conditions. Values in the "Average of curve fits" columns represent the mean ± standard error of the thermodynamic parameters obtained from individual melting curve fits. $\Delta G^{\circ}_{37}$ values were calculated from the corresponding $\Delta H^{\circ}$ and $\Delta S^{\circ}$ values at 37°C. Reported $T_M$ values are the melting temperatures calculated at 0.1 mM total oligonucleotide concentration.

| **Hairpins** | **Average of curve fits** | | | |
|---|---|---|---|---|
| | **-ΔH° (kcal/mol)** | **-ΔS° (eu)** | **$-\Delta G^{\circ}_{37}$ (kcal/mol)** | **$T_M$ (°C)** |
| GGGAUACAAAGUAUCCA | 60.4±2.8 | 175.3±8.3 | 6.02±0.26 | 71.3 |
| GGGAUACAAAAGUAUCCA | 59.3±2.8 | 170.4±8.4 | 6.43±0.18 | 74.8 |
| GGCGUAAGCC | 35.3±2.2 | 102.6±6.9 | 3.51±0.11 | 71.2 |
| GGAGUUCGCUCC | 53.5±3.6 | 158.7±11.1 | 4.25±0.32 | 63.8 |
| GGGAUACAAAAAGUAUCCA | 63.5±3.0 | 182.5±8.7 | 6.94±0.27 | 75.0 |
| GGGAUACCCCCCGUAUCCA | 45.2±1.6 | 129.7±4.7 | 4.94±0.25 | 75.1 |
| GGCAUAAUCGCC | 25.5±2.1 | 76.0±6.9 | 1.96±0.30 | 62.8 |
| GGCGUAAUAGCC | 31.8±3.7 | 93.6±11.3 | 2.74±0.28 | 66.2 |
| GGCGUAAUGGCC | 22.7±2.3 | 67.7±7.1 | 1.71±0.26 | 62.3 |
| GGCUUAAUUGCC | 28.4±1.8 | 84.7±5.8 | 2.09±0.13 | 61.7 |
| GCGAAUAAAUAUCGC | 35.0±4.7 | 105.4±14.5 | 2.27±0.30 | 58.5 |
| GCGUCCCCCCACGC | 37.0±6.9 | 112.4±21.4 | 2.14±0.28 | 56.0 |
| GGAGUAAUAUCC | 38.6±1.3 | 119.6±4.3 | 1.54±0.02 | 49.9 |
| GGUGUAAUAACC | 35.5±2.2 | 111.0±6.9 | 1.09±0.09 | 46.8 |
| GUCGUAAUAGAC | 33.6±0.6 | 104.0±2.0 | 1.37±0.04 | 50.2 |
| GGUGUAAUGACC | 33.6±1.4 | 105.8±4.2 | 0.77±0.06 | 44.3 |
| GGUUUAAUUACC | 34.4±1.2 | 109.0±3.9 | 0.53±0.05 | 41.9 |

**Supplementary Table 10. Comparison of special first mismatch bonuses for RNA hairpins.** Total hairpin free energies ($\Delta G°_{37}$) were determined experimentally by analyzing the average melting curve fits. Loop and initiation free energies were derived using the appropriate nearest-neighbor stem parameters (Adv. DMEM or 1 M NaCl) and corresponding first mismatch contributions. Special first mismatch bonus terms are shown for comparison between derived in Adv. DMEM, derived using 1 M NaCl parameters, and the theoretical values assigned by the Tuner (2004) nearest neighbor model [2,5]. For GUCGUAAUAGAC (*), the First Stack Bonus $\Delta G°_{37}$ for 1 M NaCl was determined using the corresponding value for GGCGUAAUAGCC because the original sequence had not been studied in 1 M NaCl. This substitution was considered appropriate because both hairpins contain the identical loop sequence and closing GC base pair, differing only in the adjacent stem sequence; therefore, the loop contribution was assumed to be equivalent for calculation of the derived first mismatch bonus. Turner 2004

| Sequence | Hairpin First Mismatch | Total $\Delta G°_{37}$ Adv. DMEM (kcal/mol) | Hairpin Loop $\Delta G°_{37}$ Adv. DMEM (kcal/mol) | Hairpin Initiation $\Delta G°_{37}$ Adv. DMEM (kcal/mol) | First Stack Bonus $\Delta G°_{37}$ Adv. DMEM (kcal/mol) | First Stack Bonus $\Delta G°_{37}$ 1 M NaCl (kcal/mol) | First Stack Bonus $\Delta G°_{37}$ 1 M NaCl (Turner 2004) (kcal/mol) |
|---|---|---|---|---|---|---|---|
| GGC<u>GUAA</u>GCC | CG GA | -3.51 ± 0.14 | 3.06 ± 0.67 | 4.46 ± 1.25 | -1.10 ± 1.34 | -1.29 ± 0.58 | -0.90 ± 0.10 |
| GGC<u>GUAAUA</u>GCC | CG GA | -2.74 ± 0.11 | 3.83 ± 0.67 | 5.23 ± 1.25 | -0.17 ± 1.35 | -1.07 ± 0.59 | -0.90 ± 0.10 |
| GGC<u>GUAAUG</u>GCC | CG GG | -1.71 ± 0.07 | 4.86 ± 0.66 | 6.46 ± 1.31 | 1.06 ± 1.40 | -1.34 ± 0.55 | -0.80 ± 0.30 |
| GGC<u>UUAAUU</u>GCC | CU GU | -2.09 ± 0.08 | 4.48 ± 0.66 | 5.68 ± 1.30 | 0.26 ± 1.39 | -4.07 ± 0.64 | -0.90 ± 0.10 |
| GGA<u>GUAAUA</u>UCC | AG UA | -1.54 ± 0.06 | 3.50 ± 0.61 | 4.30 ± 1.22 | -1.10 ± 1.32 | -1.46 ± 0.74 | -0.90 ± 0.10 |
| GGU<u>GUAAUA</u>ACC | UG AA | -1.09 ± 0.04 | 3.96 ± 0.63 | 5.06 ± 1.26 | -0.34 ± 1.35 | -1.08 ± 0.66 | -0.90 ± 0.10 |
| GUC<u>GUAAUA</u>GAC* | CG GA | -1.37 ± 0.05 | 2.96 ± 0.61 | 4.36 ± 1.25 | -1.02 ± 1.35 | -1.07* ± 0.59 | -0.90 ± 0.10 |
| GGU<u>GUAAUG</u>ACC | UG AG | -0.77 ± 0.03 | 4.28 ± 0.63 | 5.48 ± 1.33 | 0.03 ± 1.42 | -0.54 ± 0.69 | -0.80 ± 0.30 |
| GGU<u>UUAAUU</u>ACC | UU AU | -0.53 ± 0.02 | 4.52 ± 0.63 | 5.02 ± 1.30 | -0.37 ± 1.40 | -0.97 ± 0.67 | -0.90 ± 0.10 |

**Supplementary Table 11. Thermodynamic parameters of RNA duplexes containing bulges.** RNA duplexes containing bulges are represented by two sequences (5′–3′) for non-self-complementary helices, with dashed lines ("-") indicating gaps in alignment. The first duplex is the reference duplex, which does not contain a bulge. Thermodynamic parameters were determined by UV optical melting under Adv. DMEM conditions. Values in the "Average of curve fits" columns represent the mean ± standard error of the thermodynamic parameters obtained from individual melting curve fits. Values in the "$T_M^{-1}$ vs. log $C_T$ plots" columns were obtained from linear regression of the reciprocal melting temperature ($T_M^{-1}$) as a function of the logarithm of total strand concentration ($C_T$). $\Delta G^\circ_{37}$ values were calculated from the corresponding $\Delta H^\circ$ and $\Delta S^\circ$ values at 37°C. Reported $T_M$ values are the melting temperatures calculated at 0.1 mM total oligonucleotide concentration.

| Dupleks 5'–3' | Average of curve fits | | | | $T_M^{-1}$ vs log $C_T$ plots | | | |
|---|---|---|---|---|---|---|---|---|
| | -ΔH° (kcal/mol) | -ΔS° (eu) | -ΔG°37 (kcal/mol) | $T_M$ (°C) | -ΔH° (kcal/mol) | -ΔS° (eu) | -ΔG°37 (kcal/mol) | $T_M$ (°C) |
| CGAGGAGC<br>GCUCCUCG | 84.7±2.6 | 227.8±7.8 | 14.01±0.19 | 67.1 | 88.6±2.4 | 293.3±6.9 | 14.35±0.21 | 67.0 |
| CGAGAGAGC<br>GCUC---CUCG | 69.9±3.8 | 194.3±11.9 | 9.58±0.15 | 51.2 | 62.0±2.2 | 170.0±6.7 | 9.26±0.10 | 51.3 |
| CGAGAAGAGC<br>GCUC----CUCG | 57.8±3.6 | 160.9±11.5 | 7.88±0.10 | 44.4 | 62.3±3.6 | 175.1±11.3 | 7.99±0.09 | 44.4 |
| CGAGAAAGAGC<br>GCUC------CUCG | 59.4±12.7 | 169.7±41.7 | 6.74±0.30 | 38.1 | 51.7±2.8 | 144.8±9.0 | 6.76±0.04 | 38.4 |
| CGAGUGAGC<br>GCUC---CUCG | 70.9±5.2 | 196.8±16.2 | 9.89±0.21 | 52.4 | 66.6±2.9 | 183.3±9.0 | 9.69±0.14 | 52.5 |
| CGAGUUGAGC<br>GCUC-----CUCG | 70.0±3.1 | 198.9±9.8 | 8.31±0.12 | 45.1 | 70.6±2.9 | 201.0±9.3 | 8.31±0.05 | 45.0 |
| CGAGUUUGAGC<br>GCUC-------CUCG | 60.8±6.3 | 172.4±20.6 | 7.35±0.12 | 41.3 | 67.6±2.6 | 194.2±8.4 | 7.33±0.03 | 40.7 |
| CGAG--GAGC<br>GCUCACUCG | 57.9±11.9 | 164.7±39.2 | 6.77±0.28 | 38.3 | 51.3±3.3 | 143.4±10.5 | 6.79±0.06 | 38.6 |
| CGAG----GAGC<br>GCUCAACUCG | 56.2±5.9 | 157.4±19.1 | 7.39±0.14 | 41.8 | 58.3±1.8 | 164.4±5.9 | 7.31±0.20 | 41.2 |
| CGAG-------GAGC<br>GCUCAAACUCG | 58.5±11.5 | 166.6±37.1 | 6.84±0.15 | 38.7 | 61.6±2.1 | 176.1±6.9 | 6.92±0.02 | 39.0 |
| CGAG--GAGC<br>GCUCUCUCG | 72.1±5.0 | 201.7±15.4 | 9.49±0.20 | 50.3 | 79.6±3.3 | 225.1±10.3 | 9.74±0.10 | 50.0 |

| | | | | | | | | | |
|---|---|---|---|---|---|---|---|---|---|
| CGAG----GAGC<br>GCUCUUCUCG | 76.9±9.3 | 216.4±28.5 | 9.80±0.53 | 50.8 |  | 67.5±2.0 | 187.5±6.1 | 9.38±0.01 | 50.7 |
| CGAG-------GAGC<br>GCUCUUUCUCG | 65.6±8.2 | 186.3±26.3 | 7.83±0.14 | 43.3 |  | 64.0±3.2 | 180.8±10.2 | 7.89±0.07 | 43.7 |

**Supplementary Table 12. Thermodynamic parameters of RNA duplexes containing various internal loops.**

RNA duplexes containing various internal loops are represented by two sequences (5′–3′) for non-self-complementary helices. The first duplex is the reference duplex, which does not contain an internal loop. Thermodynamic parameters were determined by UV optical melting under Adv. DMEM conditions. Values in the "Average of curve fits" columns represent the mean ± standard error of the thermodynamic parameters obtained from individual melting curve fits. Values in the "$T_M^{-1}$ vs. log $C_T$ plots" columns were obtained from linear regression of the reciprocal melting temperature ($T_M^{-1}$) as a function of the logarithm of total strand concentration ($C_T$). $\Delta G^{\circ}_{37}$ values were calculated from the corresponding $\Delta H^{\circ}$ and $\Delta S^{\circ}$ values at 37°C. Reported $T_M$ values are the melting temperatures calculated at 0.1 mM total oligonucleotide concentration.

| **Dupleks 5'–3'** | **Average of curve fits** | | | | **$T_M^{-1}$ vs log $C_T$ plots** | | | |
|---|---|---|---|---|---|---|---|---|
| | **-ΔH° (kcal/mol)** | **-ΔS° (eu)** | **-ΔG°37 (kcal/mol)** | **$T_M$ (°C)** | **-ΔH° (kcal/mol)** | **-ΔS° (eu)** | **-ΔG°37 (kcal/mol)** | **$T_M$ (°C)** |
| CGAGGAGC<br>GCUCCUCG | 84.7±2.6 | 227.8±7.8 | 14.01±0.19 | 67.1 | 88.6±2.4 | 293.3±6.9 | 14.35±0.21 | 67.0 |
| CGAGAGAGC<br>GCUCACUCG | 77.3±8.6 | 218.2±27.4 | 9.59±0.18 | 49.8 | 78.3±2.9 | 221.2±9.0 | 9.67±0.09 | 50.0 |
| CGAGAAGAGC<br>GCUCAACUCG | 72.8±3.4 | 204.6±10.8 | 9.32±0.14 | 49.4 | 80.1±6.0 | 227.3±18.6 | 9.56±0.21 | 49.2 |
| CGAGAAAGAGC<br>GCUCAAACUCG | 71.8±4.3 | 205.5±13.7 | 8.03±0.10 | 43.6 | 76.3±1.6 | 219.9±5.1 | 8.10±0.03 | 43.5 |
| CGAGUGAGC<br>GCUCUCUCG | 87.5±4.4 | 248.0±13.7 | 10.61±0.23 | 52.2 | 77.8±5.0 | 217.8±15.6 | 10.28±0.20 | 52.7 |
| CGAGUUGAGC<br>GCUCUUCUCG | 95.7±2.6 | 273.7±8.2 | 10.81±0.11 | 51.5 | 98.7±4.6 | 283.0±14.4 | 10.92±0.17 | 51.4 |
| CGAGUUUGAGC<br>GCUCUUUCUCG | 65.6±3.7 | 187.6±11.8 | 7.43±0.14 | 41.3 | 68.8±3.7 | 197.9±12.0 | 7.46±0.06 | 41.2 |
| CGAGGAGAGC<br>GCUCAGCUCG | 70.1±4.2 | 198.7±13.6 | 8.43±0.09 | 45.6 | 74.3±1.8 | 212.4±5.7 | 8.43±0.02 | 45.1 |
| CGAGAGGAGC<br>GCUCGACUCG | 70.7±7.2 | 200.7±22.7 | 8.40±0.21 | 45.4 | 81.2±2.0 | 233.9±6.5 | 8.60±0.03 | 45.1 |
| CGAGCAGAGC<br>GCUCACCUCG | 53.2±5.2 | 146.4±16.3 | 7.77±0.18 | 44.0 | 60.9±3.4 | 170.9±11.0 | 7.87±0.05 | 44.0 |
| CGAGCUGAGC<br>GCUCUCCUCG | 91.7±3.5 | 261.7±10.8 | 10.51±0.23 | 51.1 | 104.9±7.3 | 302.7±22.7 | 10.98±0.27 | 50.7 |

| | | | | | | | | | |
|---|---|---|---|---|---|---|---|---|---|
| CGAGUCGAGC<br>GCUCCUCUCG | 81.3±1.5 | 233.8±4.9 | 8.83±0.14 | 46.0 | | 93.2±8.3 | 271.3±26.0 | 9.08±0.22 | 45.7 |
| CGAGACGAGC<br>GCUCCACUCG | 77.3±8.6 | 222.4±27.8 | 8.36±0.05 | 44.5 | | 74.8±2.8 | 214.2±9.1 | 8.33±0.03 | 44.7 |
| CGAGAAGAGC<br>GCUCAGCUCG | 74.8±2.0 | 212.0±6.4 | 9.06±0.09 | 47.8 | | 81.2±1.8 | 231.9±5.5 | 9.22±0.05 | 47.6 |
| CGAGAAGAGC<br>GCUCGGCUCG | 82.2±5.6 | 232.4±17.8 | 10.08±0.14 | 51.0 | | 75.4±5.4 | 211.3±17.0 | 9.87±0.19 | 51.4 |
| CGAGGAGAGC<br>GCUCAACUCG | 87.6±9.3 | 247.0±28.9 | 10.95±0.37 | 53.5 | | 82.8±9.8 | 232.1±30.0 | 10.77±0.50 | 53.7 |

**Supplementary Table 13. Comparison of MFE Accuracy Metrics.**
Mean sensitivity, positive predictive value (PPV), and F1 score for minimum free energy (MFE) secondary-structure predictions using 1 M NaCl and Adv. DMEM nearest-neighbor parameters for sequences in the RNAstructure Archive II benchmark dataset [11,12]. Sensitivity is the fraction of known base pairs correctly predicted, PPV is the fraction of predicted base pairs that are correct, and the F1 score is the harmonic mean of sensitivity and PPV [10,11,13-15]. Values represent the mean across sequences within each RNA family. P-values were calculated using one-tailed paired t-tests comparing the 1 M NaCl and Adv. DMEM predictions [16]. N indicates the number of sequences in each RNA family.

| RNA Family | 1 M NaCl Mean Sens | Adv. DMEM Mean Sens | P-value | 1M NaCl Mean PPV | Adv. DMEM Mean PPV | P-value | 1 M NaCl Mean F1 Score | Adv. DMEM Mean F1 Score | P-value | N |
|---|---|---|---|---|---|---|---|---|---|---|
| 5S rRNA | 65.83 | 61.59 | 0.999 | 60.47 | 59.78 | 0.745 | 62.95 | 60.58 | 0.999 | 1283 |
| 16S rRNA (Domain) | 61.22 | 59.28 | 0.687 | 53.71 | 54.67 | 0.407 | 56.87 | 56.60 | 0.574 | 84 |
| 16S rRNA (Full) | 45.14 | 42.89 | 0.661 | 38.40 | 38.66 | 0.480 | 41.41 | 40.60 | 0.669 | 22 |
| 23S rRNA (Domain) | 73.66 | 61.10 | 0.998 | 66.86 | 58.15 | 0.985 | 70.06 | 59.55 | 0.997 | 30 |
| 23S rRNA (Full) | 57.78 | 46.93 | 0.914 | 51.24 | 43.57 | 0.880 | 54.31 | 45.17 | 0.981 | 5 |
| Group 1 Intron | 56.46 | 52.52 | 0.936 | 46.15 | 45.20 | 0.636 | 50.22 | 48.03 | 0.955 | 98 |
| Group 2 Intron | 36.70 | 38.95 | 0.403 | 21.97 | 25.06 | 0.348 | 27.14 | 30.19 | 0.137 | 11 |
| RNAase P | 57.97 | 55.51 | 0.984 | 53.10 | 52.98 | 0.542 | 55.16 | 53.96 | 0.991 | 454 |
| SRP | 66.51 | 60.21 | 0.999 | 61.85 | 59.06 | 0.974 | 63.72 | 59.25 | 1.00 | 924 |
| Telomerase RNA | 59.23 | 60.44 | 0.361 | 42.54 | 45.40 | 0.160 | 49.40 | 51.74 | 0.219 | 37 |
| tmRNA | 45.89 | 43.59 | 0.992 | 41.39 | 41.44 | 0.478 | 43.21 | 42.18 | 0.966 | 462 |
| tRNA | 82.19 | 82.47 | 0.416 | 78.99 | 82.30 | 0.008 | 80.34 | 82.11 | 0.011 | 557 |

**Supplementary Table 14. Comparison of MEA Accuracy Metrics.**
Mean sensitivity, positive predictive value (PPV), and F1 score for maximum expected accuracy (MEA) secondary-structure predictions using 1 M NaCl and Adv. DMEM nearest-neighbor parameters for sequences in the RNAstructure Archive II benchmark dataset [11,12]. Sensitivity is the fraction of known base pairs correctly predicted, PPV is the fraction of predicted base pairs that are correct, and the F1 score is the harmonic mean of sensitivity and PPV [10,11,13-15]. Values represent the mean across sequences within each RNA family. P-values were calculated using one-tailed paired t-tests comparing the 1 M NaCl and Adv. DMEM predictions [16]. N indicates the number of sequences in each RNA family.

| RNA Family | 1 M NaCl Average Sens | Adv. DMEM Average Sens | P-value | 1M NaCl Average PPV | Adv. DMEM Average PPV | P-value | 1 M NaCl Average F1 Score | Adv. DMEM Average F1 Score | P-value | N |
|---|---|---|---|---|---|---|---|---|---|---|
| 5S rRNA | 63.88 | 61.98 | 0.965 | 61.17 | 62.68 | 0.068 | 62.36 | 62.13 | 0.664 | 1283 |
| 16S rRNA (Domain) | 60.25 | 58.75 | 0.879 | 55.28 | 56.37 | 0.173 | 57.17 | 56.86 | 0.605 | 84 |
| 16S rRNA (Full) | 47.47 | 45.08 | 0.892 | 43.44 | 43.29 | 0.528 | 44.83 | 44.69 | 0.530 | 22 |
| 23S rRNA (Domain) | 73.53 | 69.76 | 0.952 | 69.36 | 69.67 | 0.447 | 70.72 | 68.99 | 0.771 | 30 |
| 23S rRNA (Full) | 61.60 | 50.49 | 0.986 | 58.14 | 53.18 | 0.887 | 56.22 | 50.80 | 0.928 | 5 |
| Group 1 Intron | 56.99 | 53.15 | 0.915 | 49.37 | 48.87 | 0.565 | 52.35 | 50.34 | 0.965 | 98 |
| Group 2 Intron | 41.16 | 43.15 | 0.359 | 27.49 | 31.59 | 0.257 | 32.62 | 36.14 | 0.091 | 11 |
| RNAase P | 60.79 | 57.70 | 0.998 | 58.69 | 58.41 | 0.598 | 59.38 | 57.75 | 0.998 | 454 |
| SRP | 63.82 | 58.49 | 0.999 | 60.49 | 58.83 | 0.872 | 61.61 | 58.13 | 1.00 | 924 |
| Telomerase RNA | 58.26 | 60.38 | 0.277 | 43.45 | 46.94 | 0.120 | 49.66 | 52.71 | 0.144 | 37 |
| tmRNA | 46.03 | 43.82 | 0.991 | 44.22 | 44.85 | 0.257 | 44.78 | 43.95 | 0.938 | 462 |
| tRNA | 82.81 | 81.07 | 0.920 | 80.99 | 82.89 | 0.067 | 81.63 | 81.61 | 0.514 | 557 |

**Supplementary Table 15. Comparison of NED Scores.**
Mean NED scores for sequences within each RNA family predicted using the 1 M NaCl and Adv. DMEM nearest neighbor models. NED estimates the probability that a nucleotide is incorrectly folded relative to the known secondary structure across the thermodynamic ensemble, with lower NED values indicating greater prediction accuracy [17]. P-values were calculated using one-tailed paired *t*-tests comparing NED scores between the 1 M NaCl and Adv. DMEM models. Statistical significance was defined as $P < 0.05$. NED scores were averaged across sequences within each RNA family.

| RNA Family | 1 M NaCl NED Score | Adv. DMEM NED Score | P-value |
|---|---|---|---|
| 5S rRNA | 0.377 | 0.374 | 0.137 |
| 16S rRNA (Domain) | 0.394 | 0.381 | 0.016 |
| 16S rRNA (Full) | 0.482 | 0.493 | 0.104 |
| 23S rRNA (Domain) | 0.308 | 0.331 | 0.989 |
| 23S rRNA (Full) | 0.397 | 0.446 | 0.988 |
| Group 1 Intron | 0.426 | 0.424 | 0.362 |
| Group 2 Intron | 0.561 | 0.515 | 0.001 |
| RNAase P | 0.386 | 0.388 | 0.846 |
| SRP | 0.380 | 0.400 | 1.00 |
| Telomerase RNA | 0.451 | 0.422 | 0.014 |
| tmRNA | 0.468 | 0.461 | 0.005 |
| tRNA | 0.301 | 0.257 | $2.21\times10^{-33}$ |
